# Bidirectional allosteric ligand regulation in a central glycolytic enzyme

**DOI:** 10.64898/2026.02.05.704047

**Authors:** Belen Sundberg, Chenlin Lu, Malcolm L. Wells, Kyle C. Weber, Zhen Gong, Anum Glasgow

**Affiliations:** Department of Biochemistry and Molecular Biophysics, Columbia University, New York, NY 10032, USA

## Abstract

Allosteric regulation enables fine-tuned control of enzyme activity in response to cellular signals, yet its molecular basis often remains unclear. Phosphofructokinase-1 (PFK), the rate-limiting enzyme of glycolysis, is a paradigmatic, well-conserved system whose reaction kinetics conform to the Monod-Wyman-Changeux model of allostery. However, X-ray crystal structures of bacterial PFK orthologs in distinct ligand-bound states do not show the consistent, concerted structural rearrangements expected for classical “relaxed” and “tense” states, revealing a decades-long disconnect between structure and function. We resolve this paradox by integrating biophysical and computational approaches to show that activator and inhibitor binding to the same allosteric pocket differentially reweight the conformational ensemble of *Escherichia coli* PFK. Activator binding stabilizes conformational substates that preorganize the catalytic site, whereas inhibitor binding upweights apo-like, catalytically incompetent substates. These findings establish an ensemble-based mechanism for PFK regulation and provide an energetic framework for understanding the expanded allosteric architecture of higher PFK orthologs.

## Introduction

Allosteric regulation allows enzymes to convert molecular signals into coordinated metabolic and regulatory responses, underpinning cellular homeostasis across all domains of life. Phosphofructokinase-1 (PFK; EC 2.7.1.11) is an almost universally conserved allosteric enzyme that catalyzes the committed step of glycolysis, performing the ATP-dependent phosphorylation of fructose-6-phosphate (F6P) to generate fructose-1,6-bisphosphate (FBP).^1,2^ As a primary control point for glycolytic flux, PFK has long served as a central model for understanding the molecular basis of metabolic regulation. Bacterial PFK-1 are regulated by two primary effectors, magnesium-adenosine diphosphate (Mg-ADP) and phosphoenolpyruvate (PEP), which bind to the same allosteric site (E site) but activate or inhibit the enzyme, respectively (**Fig. 1A, B**).^3^ In contrast, eukaryotic PFK are subject to more complex combinatorial regulation, with more than ten effectors identified for human PFK, as well as regulation by post-translational modifications.^4–10^

**Figure 1.**
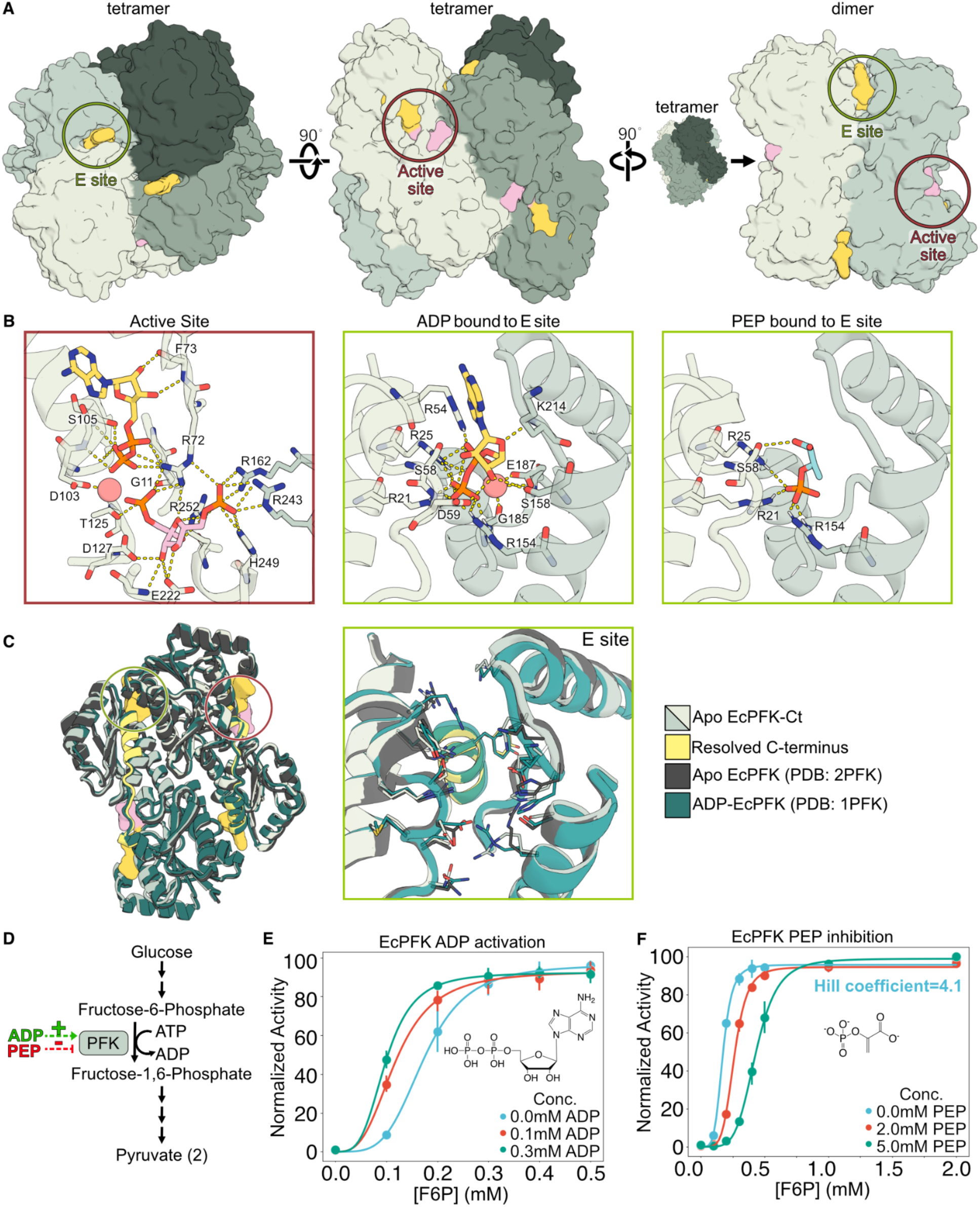
Activation and inhibition of EcPFK without large structural changes. **(A)** Newly solved X-ray crystal structure of apo EcPFK (PDB: 10OJ) in its tetramer and dimer forms, with ligand binding sites boxed. **(B)** ADP and FBP bound in the active site of the ADP-bound crystal structure (PDB: 1PFK), ADP bound in the E site in the same structure, and a structural model of EcPFK with PEP bound in the E site. Dashes denote polar contacts. **(C)** Apo EcPFK structure (PDB: 10OJ) with the newly resolved C-terminus highlighted in yellow, aligned with previously solved ADP-bound EcPFK and partial apo structures. Ligand binding sites are colored as in (A). **(D)** PFK catalyzes the rate limiting step in glycolysis. **(E, F)** Enzymatic activity assays show activation of EcPFK by ADP and inhibition by PEP. n = 3 technical replicates and 2 biological replicates were used.

In addition to catalyzing a key step in glycolysis, bacterial PFK orthologs have long been regarded as canonical systems for the Monod-Wyman-Changeux (MWC) two-state model of allostery, where effector binding shifts the equilibrium between a high-affinity, active, “relaxed” (R) state and a low-affinity, inactive, “tense” (T) state.^11–18^ Early structural studies of *Bacillus stearothermophilus* PFK (BsPFK) supported this model, revealing quaternary rearrangements, including dimer rotations and substrate-binding loop rearrangements, that provided a structural basis for the functional distinction between R- and T-states (**Fig. S1A**).^19–21^ However, even in BsPFK, ligand binding in the E sites induces minimal local structural changes, and mutational and thermodynamic analyses indicate that the crystal structures alone cannot account for the magnitude or specificity of allosteric coupling.^22–24^ Together, these findings underscore that structural differences observed in X-ray crystal structures are insufficient to explain how regulatory signals transduce over a 30 Å distance to remodel the active site across multiple protomers.

Several studies have highlighted the limitations of extending the two-state interpretation of BsPFK allosteric regulation to other bacterial PFK orthologs.^12,14,25–28^ Structural analyses of *E. coli* PFK (EcPFK) have revealed intrinsic asymmetry within a single ADP-bound R-state crystal, where protomers adopt distinct “open” and “closed” conformations.^15^ This behavior is inconsistent with the concerted, symmetric transitions proposed by the MWC model and observed in BsPFK. While effector sites differ between the apo and ADP-bound EcPFK structures, their active sites show few corresponding structural differences.^14^ Contrary to expectations that the apo EcPFK structure look like the T-state of BsPFK, it instead closely resembles the activator-bound EcPFK structure.^14^ To date, a structure of inhibitor-bound EcPFK has not been solved, further complicating the structural definition of functional states for this model system. Thus, while activators and inhibitors allosterically control PFK to tune its function in cellular metabolism, crystallographic structures alone remain insufficient to map the functional R- and T-states defined by the MWC model to distinct conformational states.

This study reconciles long-standing structural observations with the MWC framework for PFK by resolving the apparent discrepancy between structure and function. We present a generalizable experimental and computational approach that supports an ensemble-based model of regulation, in which R- and T-states emerge from ligand-dependent reweighting of a conformational ensemble rather than from discrete structural transitions.^24,26,27,29–31^ By treating PFK as a probability-weighted distribution of interconverting substates, this framework unifies the enzyme’s structural and functional behavior.

## Results

### Bidirectional ligand regulation without observable structural changes

The smallest active assembly for bacterial PFK orthologs is a tetramer, composed of a dimer of dimers.^15,21,32^ Several bacterial PFK structures have been solved, all containing four identical active sites located at the tetramer interface and four allosteric effector (E) sites at the dimer interfaces (**Fig. 1A, B**).^14,15,20,21^ Unlike eukaryotic PFKs that undergo effector-dependent changes in oligomerization between active tetramers and inactive dimers, ligand regulation of most bacterial PFKs, including EcPFK, preserves a tetrameric assembly (**Fig. S2**).^4,21,33,34^

Previous biochemical studies of EcPFK determined that, in the absence of activating ligands, the protein preferentially populates a low-affinity T-state-biased conformational ensemble.^16,35^ Consistent with this observation, X-ray crystal structures suggest that when ADP is not bound in the E site, the pocket opens and the C-terminal helix that forms part of this site becomes disordered. Accordingly, the C-terminal helix was not resolved in the EcPFK apo structure (PDB: 2PFK).^14^ However, despite clear effects on catalytic activity, ligand-dependent structural rearrangements are largely restricted to the E site. Structural changes in the active site are minimal, and the substantial quaternary rearrangements characteristic of the T- to R-state transition in BsPFK, specifically the rotation of rigid dimers within the tetramer, are absent in EcPFK (**Fig. S1**).^14^ Consistent with this, the root-mean-square deviation of alpha carbons (RMSD_Cɑ_) between ADP-bound and apo EcPFK is low (0.95 Å) (**Fig. 1C**). In any case, a detailed comparison between ADP-bound and apo state crystal structures was mired by missing electron density in the C-terminal region of the apo structure, leaving open the possibility that ligand binding may substantially restructure this helix in the E site.

To provide a more complete framework for studying allosteric ligand regulation in EcPFK, we solved the first full-length apo EcPFK structure at 2.4 Å resolution (**Fig. 1C**), which includes the C-terminal loop-helix region in the E site that was previously missing (E301-Y319). Our structure is highly similar to both the earlier apo and ADP-bound EcPFK structures (RMSD_Cɑ_=0.50 Å, 1.04 Å, respectively), underscoring the absence of major differences between the low-energy conformations captured by X-ray crystallography in the two functional states (**Fig. 1C**). Finally, we performed a coupled auxiliary enzyme assay to confirm that purified EcPFK is activated by ADP and inhibited by PEP (**Fig. 1D-F**, see Methods).^36^

### Ligands regulate EcPFK through dramatic changes in local ensemble energies

Because static X-ray crystal structures collected at cryogenic temperatures provide limited insight into the molecular basis for EcPFK ligand regulation, we used hydrogen-deuterium exchange with mass spectrometry (HX/MS) to probe how ADP and PEP binding reshape the protein’s conformational ensemble – the population-weighted set of structural substates that a protein samples – at near-physiological temperature. HX/MS reports on localized ensemble stability in proteins by measuring the time-dependent exchange of backbone amide hydrogens with deuterium in D_2_O buffer. The H-D exchange rate for each residue is determined primarily by its participation in backbone hydrogen bonds, which allows the technique to capture localized energetic changes across functional states.^37^ Although HX/MS data are traditionally limited to peptide-level analysis, recently developed methods enable the unambiguous determination of single-residue free energies of opening^38,39^, ΔG_op_, for EcPFK in three functional states (apo, ADP-bound, and PEP-bound) (**Figs. 2A, B; S3A**). Residue-level ΔG_op_ values provide a quantitative measurement of the local protein stability in the context of its conformational ensemble, and although the data themselves do not produce a protein structure, their interpretation is improved by experimentally determined atomic structures to contextualize residue-level energetic measurements. As such, differences in ΔG_op_ between states for each residue (ΔG_op, APO_-ΔG_op, HOLO_, or ΔΔG_op, APO-HOLO_) report ligand-induced changes in residue-level stability within the protein ensemble (**Fig. 2C**).

**Figure 2.**
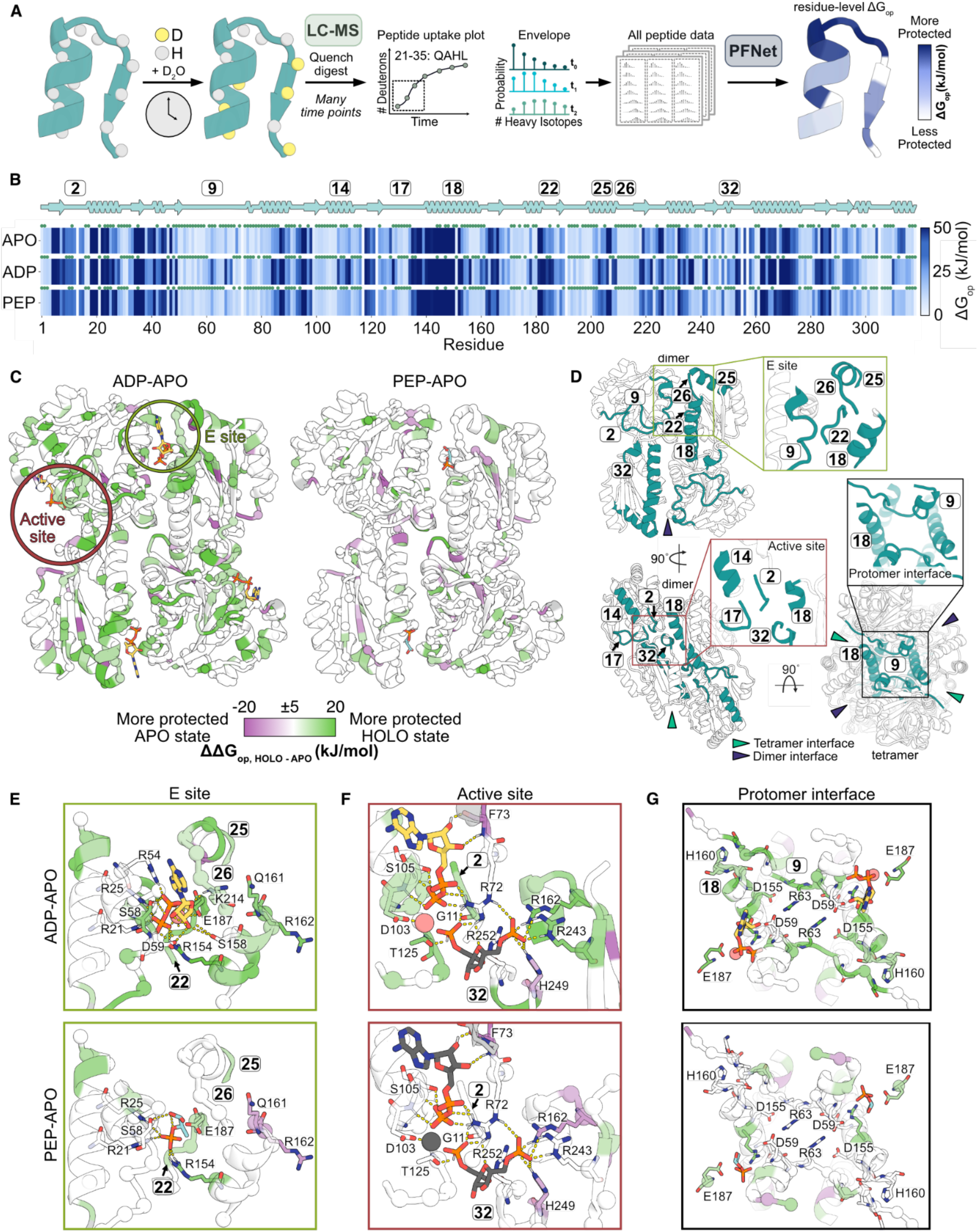
Residue-level energetic effects of ADP and PEP binding on EcPFK. **(A)** HX/MS workflow to determine residue-level ΔG_op_. **(B)** State-dependent ΔG_op_ calculated using PFNet. Darker blue indicates higher ΔG_op_, increased protection from hydrogen exchange, and increased local stability. Green circles denote residues with PFNet prediction confidence >0.8. See also Fig. S3A.**(C)** Difference in ΔG_op_ between apo and holo ensembles (ΔΔG_op, HOLO-APO_) mapped onto the crystal structure of EcPFK (PDB: 1PFK). Green residues are stabilized in the ligand-bound state relative to the apo state. Purple residues are stabilized in the apo state relative to the ligand-bound state. See also Fig. S3B, C. **(D)** Stabilized secondary structures in the ADP-apo comparison relative to the PEP-apo comparison, corresponding to regions highlighted in (E-G). **(E-F)** Insets from (C) with zoomed views of the E and active sites. Ligands are shown in gray when absent from the HX/MS experiment and are included solely to illustrate binding modes observed in the crystal structure. Residues are colored as in (C). **(G)** ΔΔG_op_ values at the protomer-protomer interfaces (top-down view), corresponding to the regions highlighted in (D).

To determine residue-level ΔG_op_ values for apo (non-liganded), ADP-bound, and PEP-bound EcPFK, we performed seven HX/MS experiments for each state, comprising three biological replicates (independently purified EcPFK samples), each with one to two technical replicates (**Table S1**). Data were pooled across experiments that incorporated three different protease conditions and 12 timepoints that spanned 30 seconds to 27 hours. The resulting dataset covered 98% of the EcPFK sequence, yielding more than 600 unique peptides per state. Following quantitative HX/MS analysis by PFNet,^38,40^ 75-78% of residues per state were resolvable to single-site ΔG_op_ values (**Figs. 2B; S3A; Table S2**). We defined high-confidence residues as those with a PFNet prediction confidence >0.8, which comprised ≥50% of all residues for each state (see Methods; **Tables S1, S2**).

HX/MS revealed that ADP and PEP binding produce distinct energetic effects on EcPFK relative to the apo ensemble. ADP binding induces widespread stabilization, reflected by elevated ΔG_op_ values across the protein, whereas PEP binding produces comparatively modest effects, resulting in a largely apo-like ensemble with more prevalent destabilization than observed upon ADP binding (**Fig. 2C**). Relative to the apo state ensemble, PEP binding induces few ligand-specific energetic effects, with changes in ΔG_op_ occurring largely at the same residues affected by ADP binding (**Figs. 2C; S3B, C**).

These contrasting energetic signatures are consistent with differences in how ligands engage the allosteric site: a prior modeling study showed that ADP binds the E site in a predominantly single, well-coordinated conformation with multiple surrounding interaction hotspots, whereas PEP exhibits greater conformational heterogeneity, consistent with its more limited effect on ensemble stabilization.^41^

As expected in direct ligand interactions, ADP binding stabilizes residues in the E site, including those on segment 9 (residues 9-15), a protomer interface loop, as well as residues in segments 18 (residues 139-160), 25 (residues 198-211), and 26 (residues 212-215), which are involved in Mg-ADP binding on the opposing dimer (**Fig. 2C, D**; segment definitions in **Table S3**).^42^ In contrast, PEP binding produces an energetic signature that largely mirrors the apo ensemble, with only limited differences in ΔG_op_ in the E site (**Figs. 2C; S3B, C**).

These distinct binding modes manifest as unique residue-specific stabilization patterns in the E site. For example, R154 is stabilized in both ligand-bound ensembles (ΔΔG_op, ADP-APO_=18.3 kJ/mol, ΔΔG_op, PEP-APO_=11.6 kJ/mol) (**Figs. 2C, E; S3C**). In the ADP-bound crystal structure (PDB: 1PFK), R154 coordinates the phosphate moiety of ADP, and its stabilization by PEP likely reflects a similar phosphate-mediated interaction. E187, which coordinates Mg^2+^ in the Mg-ADP complex, is likewise stabilized by both ligands, albeit more strongly by ADP (ΔΔG_op, ADP-APO_=12.3 kJ/mol, ΔΔG_op, PEP-APO_=6.0 kJ/mol). Despite this shared pattern, not all residues involved in ligand binding exhibit detectable backbone protection from HX. R21 and R25, which are critical for binding both ligands, show no significant backbone stabilization relative to apo in either ensemble, suggesting that their contributions to ligand binding arise primarily through sidechain-mediated interactions (**Fig. S3C**).^13,15^ Additionally, a subset of residues exhibit opposing shifts in stability depending on the bound ligand: active site residues Q161 and R162 are stabilized by ADP (ΔΔG_op, ADP-APO_=10.8, 12.9 kJ/mol, respectively), but destabilized by PEP ΔΔG_op, PEP-APO_=-7.6, -9.0 kJ/mol, respectively) (**Figs. 2F; S3B**). Notably, these residues were initially proposed to play a central role in BsPFK regulation based on early structural models, but were later shown to be nonessential for allosteric coupling.^21,22,28^

Beyond the E site, ADP binding uniquely induces long-range stabilization throughout the enzyme, consistent with an activated, R-state ensemble. In addition to stabilizing nucleotide-binding residues in the active site, ADP binding also stabilizes fructose-binding residues ∼30 Å from the E site, including Q251, T156, R243 (ΔΔG_op, ADP-APO_=17.3, 19.0, 11.7 kJ/mol, respectively), even in the absence of fructose binding (**Fig. 2F**). In contrast, PEP binding destabilizes substrate-interacting residues within the active site, including R162 and H249, which directly coordinate the phosphate group of F6P (ΔΔG_op, PEP-APO_=-8.8, -7.0 kJ/mol, respectively and ΔΔG_op, ADP-APO_=13.0, -6.7 kJ/mol, respectively) (**Fig. 2C, D, F**). The ADP-bound ensemble further exhibits residue stabilization across protomer-protomer interfaces (**Fig. 2D, G**). Because the active site is formed at these interfaces, stabilizing them may preorganize the catalytic pocket architecture, whereas PEP-associated destabilization of substrate-binding residues is consistent with reduced active site preorganization, providing a structural basis for the ligand-dependent differences in substrate affinity observed in previous studies.^16^

Together, these observations demonstrate that ADP and PEP induce distinct changes in the thermodynamic landscape of EcPFK, influencing both orthosteric and allosteric structural organization and flexibility in the absence of large-scale conformational rearrangements. We hypothesize that these ligand-specific changes in the EcPFK conformational ensemble contribute to experimentally observed differences in its K_m_.^16^

### Reweighted ensembles show molecular details underlying ligand-driven energetic effects

While HX/MS reports the energetic changes upon ligand binding, it cannot resolve the molecular interactions underlying these effects. Conversely, molecular dynamics (MD) simulations provide the necessary atomic-level detail to interpret these energetic shifts but are intrinsically limited by their faster timescales (ns to µs). To gain insight into the molecular mechanisms driving changes in residue-level ΔG_op_, we constructed Markov state models (MSMs) from extensive unbiased MD simulations of apo, ADP-bound, and PEP-bound EcPFK (30 µs per system with adaptive sampling; see Methods and **Fig. S4**). MSMs project the protein dynamics onto slow collective modes, decomposing these motions into metastable states and their interconversion rates to produce a reweighted conformational ensemble in which each conformational state is enriched according to its energy.^43–45^ By estimating the equilibrium populations of metastable states, MSMs approximate the equilibrium conformational ensemble, allowing direct, atomistic interpretation of the residue-level ΔG_op_ measured by HX/MS under equilibrium conditions.

To build the MSMs, we used secondary structure RMSD as the key molecular feature, selected based on VAMP2 score analysis and feature robustness evaluation (**Figs. S5, S6**).^46,47^ We applied time-lagged independent component analysis (TICA) for dimensionality reduction using the secondary structure RMSD as the input features, which identifies the slowest collective motions governing transitions between metastable states (see Methods; **Figs. 3A, B; S7, S8A**).

**Figure 3.**
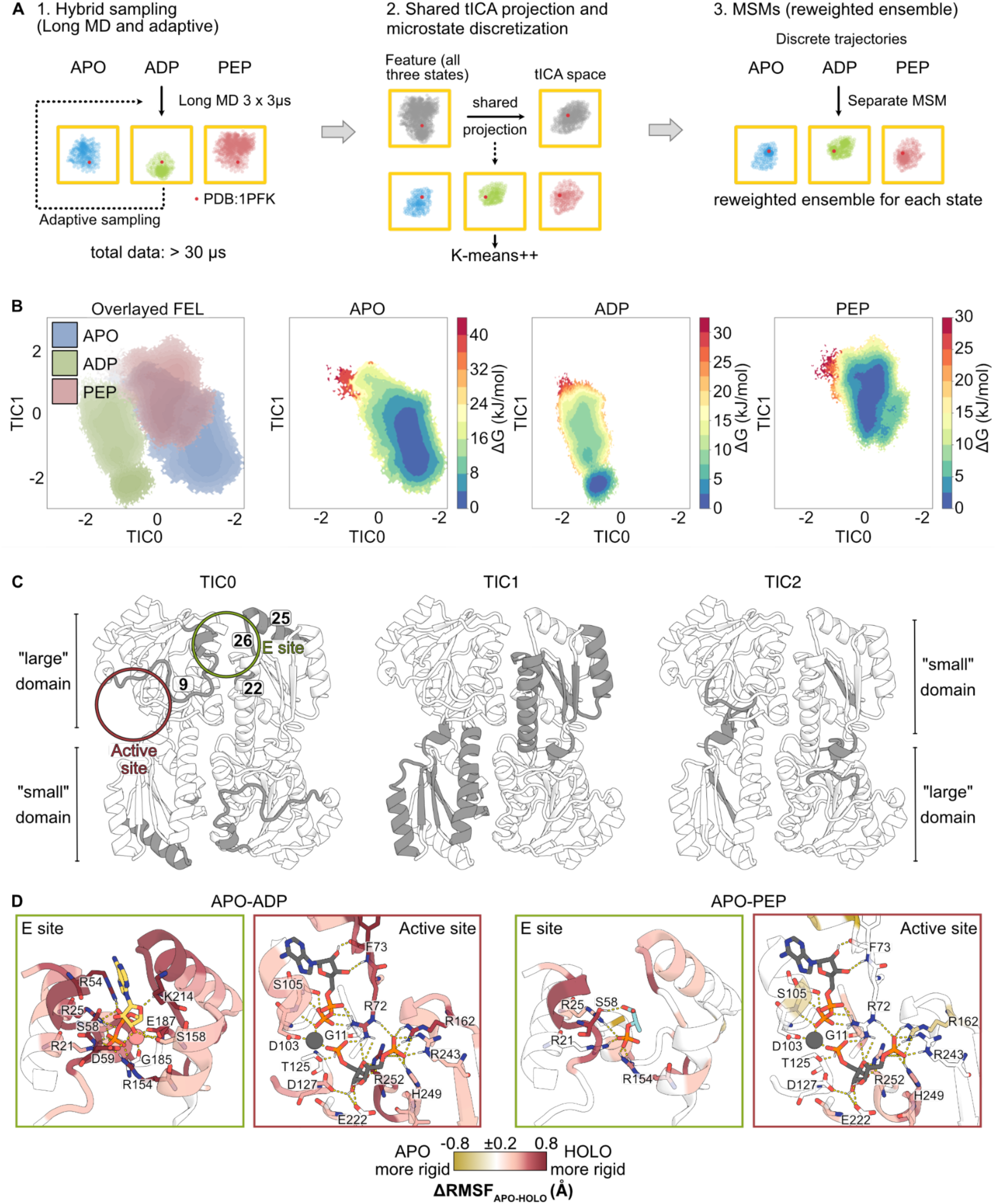
MD-derived MSMs reveal ligand-dependent redistribution of conformational states. **(A)** MD/MSM workflow. **(B)** Free energy landscape of apo, ADP-bound, and PEP-bound ensembles projected onto TIC0 vs. TIC1. The leftmost panel shows the overlay of all three ensembles, while the subsequent panels show each ensemble individually, colored by population density (ΔG). See also Figures S4-S8. **(C)** Secondary structure elements most strongly correlated with the top 3 TICs (Pearson r > 0.75) with location of ligand binding sites circled. **(D)** Difference in residue-level RMSF between apo and holo ensembles (ΔRMSF_APO-HOLO_), calculated from MSMs and mapped onto the ADP-bound EcPFK crystal structure (PDB: 1PFK). Red residues are more rigid in the holo ensemble; gold residues are more rigid in the apo ensemble. The E and active sites are stabilized in the ADP-bound ensemble relative to the apo state, with weaker stabilization observed for the PEP-bound ensemble. Ligands are shown in gray if they are absent from the MD/MSM and are included in the figure to illustrate the binding mode observed in the crystal structure.

Echoing our HX/MS results, our TICA/MSM analysis showed that ADP binding distinctly reshapes the conformational ensemble, most clearly resolved in the TIC0-TIC1 projection, while the PEP and apo ensembles sample overlapping regions in the free energy landscape (FEL). In this projection, TIC0 cleanly separates the ADP-bound ensemble from both APO and PEP ensembles, indicating that ADP binding stabilizes conformations that are largely inaccessible in the absence of an activator (**Fig. 3B**).

To identify the structural basis for this effect, we examined features most strongly correlated to the dominant collective motions in our MSMs. These features map to distinct regions of the protein, with TIC0- and TIC1-correlated secondary structures localized to the allosteric (E) site and the fructose-binding “small” domain, respectively, rather than the ATP-binding “large” domain as defined by Shirakihara and Evans (**Fig. 3C**).^15^ This spatial organization suggests ADP-induced correlated motions that couple the E site to the small domain.

Consistent with an ADP-specific role, TIC0 uniquely isolates the ADP-bound ensemble and is dominated by E site and interprotomer regions, including segment 9 (residues 9-15) at the protomer interface, and segments 22, 25, and 26, involved in Mg-ADP binding. Segments 25 (residues 198-211) and 26 (residues 212-215) form a continuous structural element at the dimer interface that contacts the ADP ribose and adenine ring moieties and may promote tighter protomer association upon ADP binding. Reinforcing this role, in our HX/MS experiments, these segments are stabilized upon ADP binding relative to the apo state, whereas PEP binding produces no detectable change, reflecting its lack of direct contacts with these regions (**Fig. 2C, E**). Segment 26 shows positive ΔΔG_op, ADP-PEP_, supporting preferential engagement of TIC0-associated segments by ADP (**Fig. S8B, C**). By contrast, segment 22 (residues 185-187), which coordinates Mg^2+^, is stabilized in both ADP- and PEP-bound states relative to apo, consistent with shared phosphate-mediated interactions rather than ADP-specific interprotomer coupling (**Fig. 2E**). Together, these observations indicate that TIC0 primarily captures the activating allosteric signal initiated by ADP binding in the E site.

TIC1 captures a complementary mode that partially separates the three ensembles, with the ADP-bound ensemble largely overlapping with the apo state ensemble and a small PEP-specific population along this coordinate (**Fig. 3B**). While TIC1 does not fully resolve ADP-specific conformations, joint projection with TIC2 reveals an ADP-specific low-energy basin that is inaccessible to both apo and PEP ensembles (**Fig. S8A**). TIC1-correlated segments map to the fructose-binding “small” domain, consistent with coordinated motions that modulate substrate-binding elements (**Fig. 3C**). Accordingly, although the apo and PEP FELs overlap along TIC0, their partial separation along TIC1 suggests that this mode distinguishes inhibitory from unliganded active site configurations, while higher-dimensional projections expose selective stabilization of catalytically activated conformations in the ADP-bound ensemble (**Fig. S8A**).

Together, this hierarchy of collective motions defines a layered allosteric mechanism in which ADP binding stabilizes unique activator-specific E site conformations (TIC0) and reweights active site and interprotomer rearrangements toward catalytically competent substates (TIC1). The energetic coupling between the E site and the active site observed in our MD-MSM analysis independently corroborates HX/MS-derived, state-specific changes in ΔG_op_ in these sites, demonstrating that these orthogonal techniques converge on a common underlying ensemble-reweighting mechanism.

### Residue-specific ensemble reweighting reveals local structural differences

To link global ensemble reweighting to local EcPFK dynamics, we computed per-residue heavy-atom root mean square fluctuation (RMSF) from the reweighted ensembles for each functional state of EcPFK, defined as the average deviation of each of a residue’s heavy atoms from their mean positions. Collectively, the average RMSF significantly differed between states (Welch’s *t*-test: APO vs. ADP, *p*=2.70×10⁻^30^; APO vs. PEP, *p*=3.98×10⁻^4^; ADP vs. PEP, *p*=4.44×10^-16^), with the largest overall reduction in flexibility observed in the ADP ensemble (**Fig. S9A**), in line with the MSM and HX/MS analyses.

Per residue ΔRMSF_APO-HOLO_ revealed that ADP binding produces widespread stabilization spanning the E site, active site, and protomer interfaces, mirroring HX/MS protection patterns (**Figs. 3D; S9B-D**). In contrast, PEP binding stabilizes fewer E and active site residues, consistent with the ΔΔG_op_ values derived from HX/MS analysis (**Figs. 2C; S3B, C**).

Although HX/MS experiments with ADP (a reaction product and allosteric activator) report on both active site and E site binding, the MD/MSM approach isolates allosteric effects arising solely from E site occupancy. Even under these conditions, ADP-dependent stabilization is clear in the E site as well as the nucleotide- and fructose-binding regions of the active site, suggesting that E site binding alone is sufficient to stabilize the catalytic region across a 30 Å distance (**Figs. 3D; S9C**). As in the HX/MS experiments, control simulations with ADP bound in both the E and active sites show similar active site stabilization (**Fig. S9E, F**).

### Biochemically diverse interactions enable global energetic changes

EcPFK undergoes pronounced ligand-dependent changes in local stability without exhibiting large-scale global conformational rearrangements, maintaining a similar overall structure across functional states. To place these local changes in the context of global ensemble behavior, we catalogued ligand-dependent rearrangements in hydrogen bonds, salt bridges, and hydrophobic interactions within individual protomers and across protomer interfaces for the reweighted ensembles (**Fig. S10**, see definitions in Methods). In agreement with TICA analysis, these rearrangements are concentrated at the E site, the protomer interfaces, and the active site.

At the E site, ADP and PEP form distinct hydrogen bonding networks. Both ligands form similar contacts with R21, R25, R154, S58 and D59. However, ADP engages with higher occupancy (61%, 65%, 79%, 74% and 90% for ADP vs. 33%, 38%, 48%, 24% and 37% for PEP) (**Fig. 4A**) and additionally forms state-specific inter-protomer contacts with H215 (17%) and K214 (60%), spanning the dimer interface. PEP instead forms state-specific contacts with K213 (14%) (**Fig. 4A**), indicating that ADP establishes a more extensive E site network with more efficient coupling across subunits.

**Figure 4.**
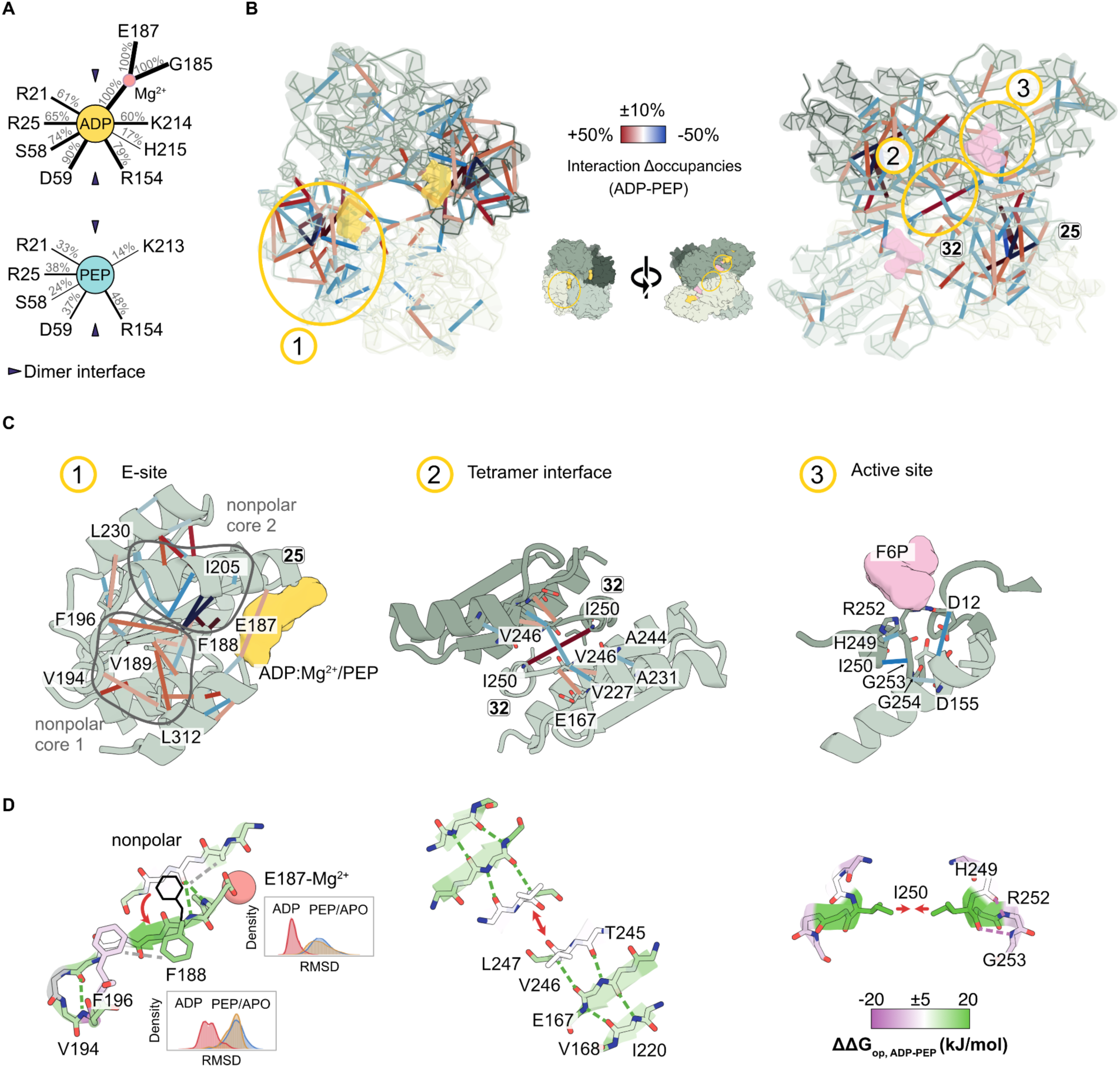
Interaction rearrangements underlying allosteric regulation. (**A**) Schematic illustrating interaction differences between the ADP- and PEP-bound ensembles at the E site. (**B**) Ligand-dependent interaction rearrangements between the ADP- and PEP-bound ensembles. Colored lines indicate changes in interaction occupancy. Hydrogen bonds, nonpolar contacts, and salt bridges are shown simultaneously; individual interaction types are shown separately in Figure S10. (**C**) Zoomed views of interactions at the E site, tetramer interface, and the active site. (**D**) ΔΔG_op, ADP–PEP_ values mapped onto the same regions shown in (C), highlighting molecular structural details.

Remarkably, these ligand-specific binding modes produce distinct interaction networks that stabilize distinct hydrophobic cores. In the ADP-bound ensemble, hydrophobic interactions F188-F196, V189-L312, V189-A181, and V194-L312 are selectively strengthened relative to PEP (Δocc_ADP-PEP_=22%, 24%, 18%, 12%, respectively), forming a nonpolar interaction core flanked by segments 25 and 40 (**Figs. 4B, C, 5B**). By contrast, the PEP-bound ensemble preferentially stabilizes an alternative hydrophobic core, flanked by segments 25 and 29, comprising enhanced F188-I205, F188-A216, V218-I163, V218-V165, and V165-L230 interactions (Δocc_ADP-PEP_=-55%, -54%, -26% -18%, -16%, respectively) (**Figs. 4B, C, 5B**).

**Figure 5.**
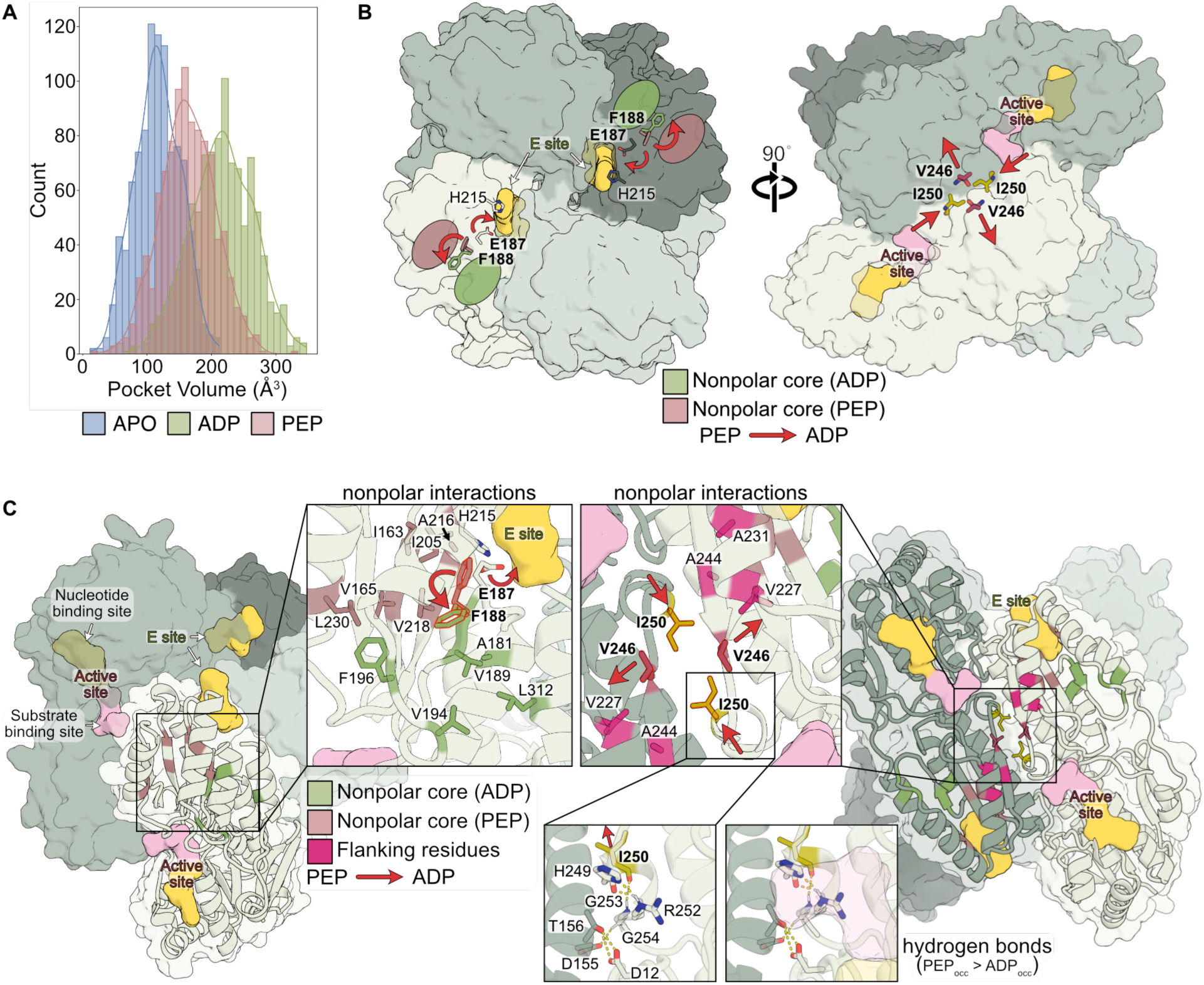
Consolidated model for EcPFK ligand regulation. **(A)** Distribution of active site pocket volumes sampled in each ensemble. See also Figure S13. **(B)** EcPFK tetramer with key residues comprising the nonpolar cores stabilized in the ADP- and PEP-bound ensembles, with flanking residues. Hydrogen bonds near the active site with higher occupancy in the PEP-bound ensemble relative to the ADP-bound ensemble are shown. Red arrows show ligand-dependent conformational changes corresponding to the transition between the PEP- and ADP-bound ensembles. **(C)** Consolidated model of EcPFK allosteric regulation by ADP and PEP, illustrating the ligand-dependent reweighting of conformational states within the tetrameric ensemble. Bolded residues in (C) and (D) denote key molecular switch residues.

These ligand-specific hydrophobic networks converge on F188, which functions as a central switch between the two cores. In the PEP-bound ensemble, the absence of Mg-ADP coordination by E187 favors an alternative F188 rotamer, biasing the enzyme toward the PEP-associated hydrophobic network (**Fig. 4D**, left). This local reorganization is reflected by distinct E187 and F188 conformations in the ADP versus PEP and APO ensembles, and positive ΔΔG_op, ADP–PEP_ HX/MS values (**Fig. 4D**, left). Notably, both F188 rotamers are present in existing crystal structures (**Fig. S11**), though this heterogeneity was not previously discussed. Together, these data corroborate our hypothesis that ligand-specific coordination at the E site organizes alternative hydrophobic cores.

In addition to these core rearrangements, we observed ligand-dependent switching at the tetramer interface. Interactions involving V246-V246 and I250-I250, supported by flanking interactions V246-V227 and A231-A244, respond oppositely to ADP and PEP binding (**Fig. 4C**, center). In the ADP-bound ensemble, V246-V246 contacts are weakened (Δocc_ADP-PEP_=-17%), whereas I250-I250 interactions are strengthened (Δocc _ADP-PEP_=38%), drawing protomers closer together on average. Consistent with this, HX/MS analysis shows positive ΔΔG_op, ADP–PEP_ for neighboring beta strand residues (L247, E167, and I220) (**Fig. 4D**, center), indicating strengthened backbone hydrogen bonding that disfavors the conformational flexibility required to maintain V246-V246 packing in the ADP ensemble.

These interface rearrangements are accompanied by reduced hydrogen bond occupancy in the ADP-bound ensemble at the active site. When EcPFK binds Mg-ADP in the E site, hydrogen bond occupancy is reduced for H249-R252 and I250-G253 on segment 32 in the active site within the same protomer (Δocc_ADP-PEP_=-18% and -29%, respectively), as well as G254-D155 and D12-T156 between subunits at the tetramer interface (Δocc _ADP-PEP_=-11% and -24%, respectively) (**Fig. 4C**, right). Reduced occupancy of these interactions suggests an R-state-like architecture that more frequently samples substates with an open, substrate-accessible pocket in the presence of ADP. In contrast, retention of the H249-R252 hydrogen bond in the PEP ensemble occludes the substrate-binding site, producing a steric clash between R252 and aligned FBP, in line with a suboptimal substrate-binding geometry (**Fig. S12**).

In summary, ADP and PEP binding upweight conformational states with distinct hydrophobic cores through E site-centered coordination, discrete molecular switches at F188 and the tetramer interface, and differential coupling to active site residues. These coordinated local rearrangements reshape the EcPFK conformational ensemble and promote activation vs. inhibition without requiring large-scale structural transitions.

### Ligand binding toggles active site volume

Biochemical studies have shown that ADP and PEP exert opposing effects on EcPFK substrate affinity at the active site, yet the molecular basis of this regulation remains unclear given the minimal differences between apo- and ADP-bound crystal structures.^14,15^ To test whether ligand-dependent, local rearrangements restructure the active site, we examined changes in active site volume upon ADP and PEP binding in the MSM-reweighted ensembles. Pocket volume analysis of the ensembles showed that the ADP-bound ensemble adopts the largest active site volume on average (**Figs. 5A; S13A**).

Supporting this observation, RMSD distributions of active site segments showed that ADP stabilizes a more restricted conformational arrangement of segments 2 and 32 (**Fig. S13B, C**), whereas the apo and PEP-bound ensembles display broader, overlapping distributions, with segment 32 sampling a wider range of conformations. In agreement with this computational result, HX/MS revealed ADP-dependent backbone stabilization of key active site residues within segments 2 (*e.g.*, S9, G10, G11) and 32 (*e.g.*, H249, I250) (**Fig. 2F**). Thus, while the active site adopts a similar conformation in the lowest-energy crystal structures, the underlying energetic ensembles differ considerably, with the apo and PEP-bound ensembles sampling a broader distribution of higher-energy conformations. In sum, these results indicate that ADP binding stabilizes a well-defined and more open active site configuration, providing a structural basis for the increased substrate affinity originally described by Blangy, Buc, and Monod in 1968.^16^

### Consolidated model for effector regulation of EcPFK

Our experimental and computational results support a model in which ADP and PEP regulate EcPFK not by inducing dramatic changes in oligomerization state or domain structure, but by selectively reshaping the conformational ensemble of the protein. This conformational reweighting occurs via modulation of specific inter-residue interactions within and across protomers. Central to this mechanism are distinct interaction networks, governed by residue switches, that bias the ensemble toward either catalytically competent or incompetent substates without global structural rearrangement (**Fig. 5B, C**).

ADP binding upweights conformational states that are not obvious when comparing EcPFK apo and ADP-bound crystal structures, including our new full-length apo state structure. ADP binding selectively stabilizes a subset of pre-existing conformations characterized by increased stabilization extending from the E site, across protomer interfaces, and into the active site, reflecting long-range energetic reorganization. These stabilized interactions define a distinct low-energy ensemble that is largely inaccessible in the apo and PEP-bound states (**Fig. 3B**).

At the molecular level, ADP binding favors ligand-specific hydrogen bonds at the E site that bridge the two protomers at the dimer interface, with the conformational change occurring via a F188-centered nonpolar interaction core. Mg-ADP binding additionally stabilizes distinct E187 and F188 rotamers, such that both residues act as ligand-dependent switches to bias the protein towards the ADP or PEP functional states. This network is coupled to molecular switches at V246 and I250, favoring strengthened I250-I250 and weakened V246-V246 interactions in the ADP-bound ensemble, while selectively weakening specific active site interactions to ultimately increase substrate accessibility and pocket volume.

### Two-lobed architecture of human PFK preserves bacterial allosteric behavior

Eukaryotic PFK evolved through gene duplication and tandem fusion of a bacterial-like ancestor, producing a larger tetrameric enzyme composed of a conserved N-terminal lobe and a more divergent C-terminal lobe (**Fig. 6A, B**).^2,48,49^ This expansion provides a natural framework for assessing how bacterial mechanisms of ligand-induced conformational reweighting and inter-subunit signal transduction are retained and elaborated in eukaryotic PFK.

**Figure 6.**
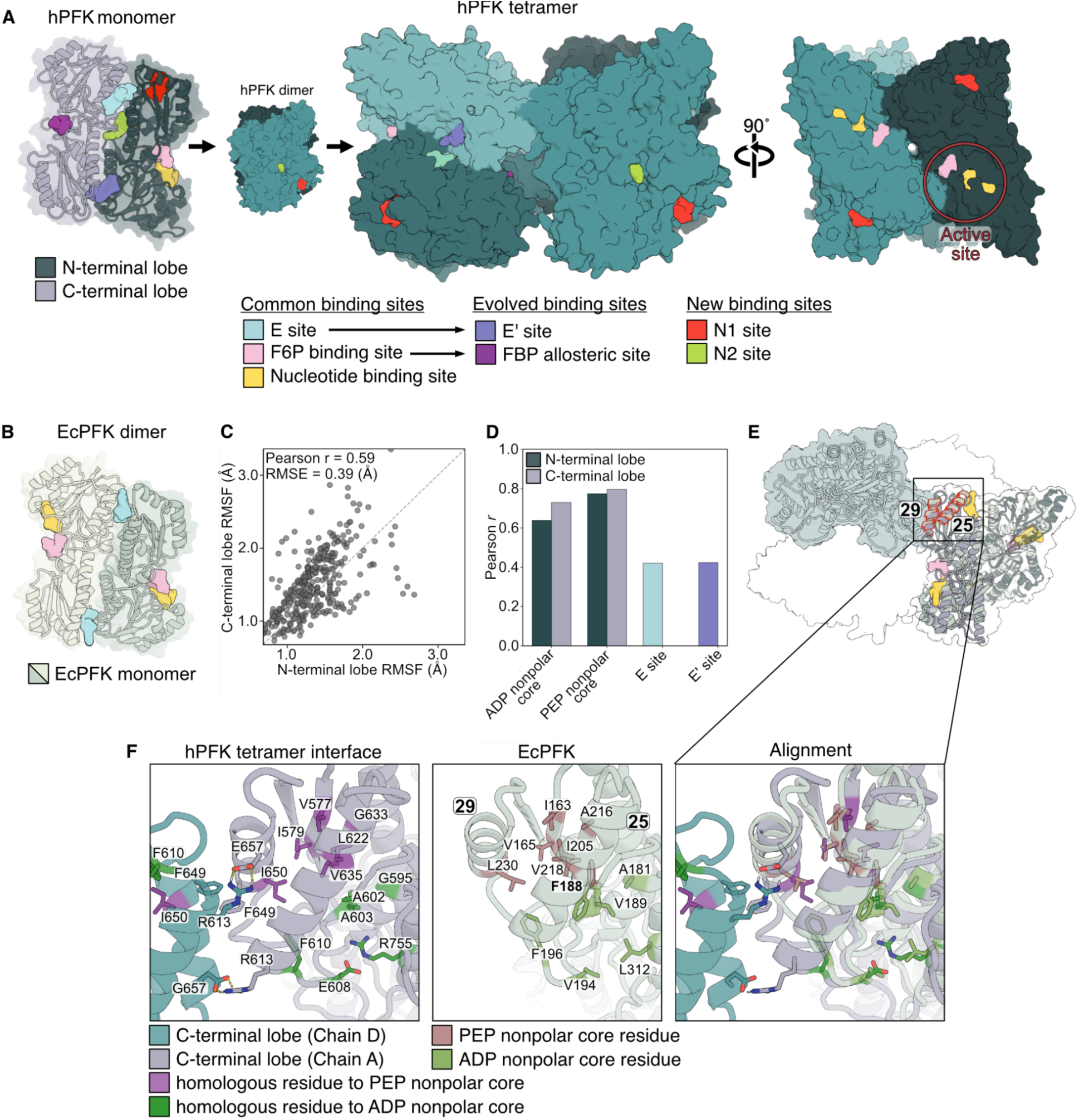
Conserved dynamical signature of bacterial PFK found in human PFK. **(A)** X-ray crystal structure of human PFK (hPFK) in tetramer, dimer, and monomer forms (PDB: 4XYJ), with ligand binding sites indicated by color. **(B)** X-ray crystal structure of the EcPFK dimer with binding sites illustrated (PDB: 1PFK), with ligand binding sites colored using the same scheme as in (A). **(C)** Scatter plot of heavy-atom RSMF for aligned residues of the hPFK N-terminal lobe vs. C-terminal lobe (n = 337). The dashed line denotes the linear relationship. See also Figure S14E. **(D)** Pearson correlation of aligned per-residue RMSF values between EcPFK and each hPFK lobe across key functional regions. The E site and E’ site are shown in distinct colors as each includes residues from both N- and C-terminal lobes. Higher Pearson r values indicate covariation in residue flexibility between aligned EcPFK and hPFK residues. See Table S4 for definitions of each site. See also Figure S14D. **(E)** Aligned structure of tetrameric hPFK and dimeric EcPFK. **(F)** Zoomed views of the hPFK tetramer interface highlighting residues homologous to those forming the ADP- and PEP-binding-induced nonpolar cores in EcPFK. Homologous residues were identified by sequence alignment of EcPFK with the N- and C-terminal lobes of hPFK by US-align.

Despite low sequence identity between the two lobes (27%) and only moderate identity to EcPFK (36% and 27% for the N- and C-terminal lobes, respectively), structural analysis of human platelet PFK (hPFK; PDB: 4XYJ) shows that the lobes adopt highly similar folds and closely resemble the EcPFK monomer, consistent with retention of a shared ancestral architecture (RMSD_Nt:Ct_=1.90 Å; RMSD_Nt:1PFK_=1.54 Å; RMSD_Ct:1PFK_=1.68 Å) (**Fig. S14A, B**).^4^ The emergent two-lobed architecture introduced additional allosteric binding sites and inter-subunit interfaces, enabling regulation by new ligands – many of them glycolysis-related metabolites – and oligomerization-dependent catalysis. Despite this increased complexity, regulation by ADP and PEP is conserved in eukaryotic PFK orthologs. Structural and mutational studies indicate that two bacterial E sites were repurposed as C-terminal sites that bind PEP and citrate (E’ sites), whereas the remaining N-terminal E sites, together with newly evolved nucleotide-binding sites, may mediate regulation by ATP and ADP (**Fig. 6A, B**).^4,50^

To determine whether the dynamic signatures of bacterial PFK are conserved in higher organisms, we performed ultra-long atomistic MD simulations of full-length, tetrameric apo hPFK using the Anton2 supercomputer (see Methods).^51^ We performed RMSF analysis of each lobe to assess inter-lobe dynamic coupling within hPFK, revealing reduced flexibility of the “modern” C-terminal lobe relative to the “ancient” N-terminal lobe and moderate correlation between aligned residues (Pearson *r* = 0.59), consistent with partial inter-lobe coupling (**Figs. 6C; S14C, E**).^52^ To assess conservation of regulatory dynamics, we compared per-residue RMSF values between apo EcPFK and hPFK across aligned residues. Consistent with greater structural and sequence similarity, RMSF correlations with EcPFK are higher for the N-terminal than the C-terminal lobe overall (Pearson *r* = 0.54 vs. 0.37) (**Fig. S14F, G**). In contrast, RMSF values of EcPFK E site residues correlate similarly with homologous residues in the E and E’ sites, which span both lobes (Pearson *r* = 0.42 for both), consistent with the retention of bacterial E site-like dynamics in the C-terminal lobe under shared PEP-binding constraints (**Figs. 6D, S14D, Table S4**). Residues comprising the ADP- and PEP-stabilized nonpolar cores have moderate to strong correlations across both lobes (Pearson *r* = 0.64 and 0.73 vs. 0.77 and 0.80) (**Figs. 6D, E, S14D, Table S4**). Together, these results suggest that the apo hPFK ensemble may preserve bacterial-like inhibitory dynamics at the E’ site and associated nonpolar interaction networks, a latent scaffold for effector regulation despite architectural divergence.

## Discussion

Phosphofructokinase was among the first enzymes described by the MWC concerted model of allostery in 1968, with early crystallographic studies of BsPFK identifying quaternary rearrangements that provided a structural basis for distinct T- and R-states.^11,16,20,21^ However, subsequent studies showed that these structural changes are insufficient to fully explain allosteric coupling between the E and active site.^22–24^ In EcPFK, this discrepancy is more pronounced: the apo and activator-bound structures are highly similar; the active site geometry remains largely conserved; the activator-bound structure exhibits structural asymmetry; and the rigid-body rearrangements characteristic of BsPFK are absent.^14,15,25,26^ Together, these findings challenged a literal interpretation of the MWC model for EcPFK and raised the long-standing question: how does EcPFK achieve bidirectional allosteric regulation in the apparent absence of conformational change?

By integrating high-resolution HX/MS experiments with X-ray crystallography, atomistic MD simulations, and Markov state models, we resolved this paradox, demonstrating that EcPFK allosteric regulation arises from ligand-dependent reweighting of the protein’s conformational ensemble, rather than transitions between discrete, homogenous structures. We found that ADP and PEP do not impose new oligomeric states or large-scale structural rearrangements on EcPFK. Instead, they bias pre-existing populations of conformational substates through selective stabilization or destabilization of specific interaction networks, invisibly reshaping the FEL. Together with more recent biophysical studies, our results support an emerging ensemble-based understanding of PFK allostery. For example, a frequency-domain fluorescence study of EcPFK demonstrated that effectors modulate the protein’s dynamics to change conformational substate populations without inducing large structural rearrangements, while NMR analyses of BsPFK revealed distributed, ligand-dependent perturbations across the protein rather than a single concerted pathway.^24,27^ Beyond PFK, the integrative, generalizable strategy presented in this study provides a means of resolving apparent discrepancies between structural and functional descriptions of allostery in other systems.

Our combined HX/MS and MD/MSM analyses revealed a striking energetic asymmetry between activation and inhibition. PEP binding largely preserves an apo-like conformational ensemble with increased sampling of catalytically inactive conformational substates, whereas ADP binding reweights the ensemble toward a low-energy subset of catalytically competent substates that are rarely sampled in the absence of an activator.

This ensemble-based mechanism reconciles a large body of prior mutational data. Residues identified by our HX/MS and MD/MSM analyses map directly onto sites previously shown to alter effector responses. In particular, E187, which we independently identify here as a ligand-dependent switch residue that distinguishes activating from inhibitory inter-residue coupling, has been shown to convert PEP into an activator when mutated to alanine.^12^ Prior studies concluded that this mutation does not impair PEP binding but instead disrupts inhibitory coupling, a finding that is fully consistent with our observation that increased occupancy of specific E187 rotamers is required for PEP to stabilize the inhibitory ensemble.^13,27,53^ Similarly, mutations at H249 and R252 selectively disrupt PEP inhibition, consistent with our observation that the H249-R252 hydrogen bond is preferentially occupied in the inhibited ensemble.^54^

More broadly, this work provides a modern reinterpretation of the MWC framework by offering a quantitative molecular mechanism for allosteric regulation that does not depend on concerted transitions between discrete R- and T-state conformations – transitions that have eluded structural characterization in EcPFK despite crystallographic studies dating back to the 1980s. Instead, EcPFK operates through ligand-dependent redistribution of populations within a largely shared conformational landscape. This mechanism inherently accommodates opposing effectors acting at the same site, as well as partial ligand occupancy and mixed regulatory states, which likely occur under physiological conditions. The ensemble-based approach also offers a powerful lens for understanding the evolution and regulation of higher PFK orthologs and a springboard for understanding pathological dysregulation of hPFK in metabolic diseases^55^ and cancers^56,57^, where disease-associated mutations are distributed throughout the protein and cannot be readily explained by models based on lowest-energy structures alone. Therapeutic development efforts may then exploit the ensemble model to precisely stabilize conformational substates associated with increased or decreased activity of this essential enzyme, thereby returning the system to homeostasis by tuning its activity allosterically and specifically rather than completely inhibiting it.

### Limitations of the study

Although this work focuses on equilibrium state ensembles, there are timescale differences between protein conformational ensembles measured by HX/MS (s-h) and MD simulations (ns-µs) that limit their direct, quantitative comparison. Constructing MSMs partially addresses this limitation by stretching the timescale of simulations to the µs-ms range, given sufficient sampling. Although we performed unbiased atomistic MD simulations extending to the tens of µs range per functional state of EcPFK and conducted appropriate validation tests in building the MSMs (**Fig. S7**), further sampling could reveal rare, high-energy conformational states not captured here. Additionally, we note that ADP binds to the E and active sites as the catalytic product of PFK and an allosteric activator. We assume that this occurred in our HX/MS experiments. To separate the effects of ADP binding in the two sites, we performed MD simulations of EcPFK with ADP bound at both sites and individual sites (**Fig. S9E, F**).

## Resource availability

### Lead contact

Requests for further information and resources should be directed to and will be fulfilled by the lead contact, Anum Glasgow.

## Materials availability

The EcPFK expression plasmid generated in this study (**Table S5**) and the *E. coli* expression strain are available from the corresponding author upon request.

## Data and code availability

- The X-ray crystallography data and refined structural models have been deposited in the Protein Data Bank under accession code 10OJ.
- Raw X-ray crystallography data, processed HX/MS peptide-level data in HXMS format^58^ and PFNet^40^ outputs, and PyMOL sessions to visualize these data, as well as molecular dynamics trajectories generated in this study, have been deposited on Zenodo at https://zenodo.org/records/18390796.
- Raw HX/MS data have been deposited to the ProteomeXchange Consortium via the PRIDE partner repository with the dataset identifier PXD073975.
- This paper does not report original code.
- Any additional information required to analyze the data reported in this paper is available from the lead contact upon request.

## Acknowledgements

The authors thank the Glasgow Lab, Dr. Bradley Webb (West Virginia University) and Dr. Liang Tong (Columbia University) for useful discussions. Dr. Wayne Hendrickson for advice on X-ray crystallography and Dr. Rinat Abzalimov at the Advanced Science Research Center at the City University of New York in his role as the MS Facility Manager. This work was supported by an ACS Catalyst Award and NIH grants R00GM135529 and R35GM157185 to AG. Anton2 computer time was provided by the Pittsburgh Supercomputing Center (PSC) through NIH grant R01GM116961. The Anton 2 machine at PSC was made available by D.E. Shaw Research. The authors also thank the staff at NYX beamline at NSLS-II for their support during data collection. BS was supported by an NIH F31 fellowship. KW was supported by an NSF Graduate Research Fellowship.

## Author contributions

BS and CL prepared the protein constructs, purified the proteins, and performed *in vitro* assays (activity assays, mass photometry, native gel electrophoresis, and SDS-PAGE). KW and ZG prepared crystallization trays, collected X-ray diffraction data, and processed crystallography datasets. KW, ZG, and CL built models. CL performed MD simulations, including adaptive sampling and MSM construction. CL and BS analyzed MD simulation data. BS and CL performed HX/MS experiments with the help of MLW. BS and CL processed and analyzed HX/MS data. BS and CL performed statistical analyses and data visualization for all experiments and computational results. AG conceived the study and supervised the research. BS, CL, and AG wrote the manuscript with input from all authors.

## METHODS

### Bacterial strains

*Escherichia coli* DH10B was used for cloning and *Escherichia coli* (*E. coli*) LOBSTR cells were used for protein expression.^59^ Cells were grown in Terrific Broth (TB) (Invitrogen) at 37 °C unless otherwise noted.

### Molecular cloning

The *E. coli* PFKA gene (GenBank: X02519.1) was cloned out of DH10B by polymerase chain reaction (PCR) using forward primer (5’-AGTATCGGTCTCCCATGCATCACCATCACCATCACGAAAACTTATATTTCCAATCTATTAAGAAAATCGGTGTGTTGACAAGCGGCG-3’) and reverse primer (5’-AGTATCGGTCTCACTTAATACATTTTTTCGGCGCAGTCCAGCCAG-3’) to insert a 6**×**His tag and TEV cleavage site at the 5’ end. This gene product was cloned into a custom pET9a vector via Golden Gate cloning using BsaI and the gene sequence was confirmed by Sanger sequencing.^60^ PCR amplification and Golden Gate cloning were performed using standard molecular biology techniques.

### Protein expression and purification

The pET9a-6×His-TEV-EcPFK plasmid was chemically transformed into *E. coli* LOBSTR cells.^59^ A single colony was grown at 37 °C in Terrific Broth (TB) (Invitrogen) containing 50 µg/mL kanamycin (ThermoFisher) for 12-16 hours overnight. The overnight culture was subcultured at a 1:50 dilution in 1 L TB containing 50 µL/mL kanamycin and grown at 37 °C while shaking at 220 rpm until the optical density at 600 nm (OD600) of the culture was between 0.4 and 0.6. The temperature was then decreased to 16 °C and expression of the target gene was induced by adding 1 mM IPTG (GoldBio) and growing the culture for an additional 12 h with shaking at 220 rpm. The culture was centrifuged at 6000 × *g* for 30 minutes at 4 °C and the pellet was frozen at -20 °C for at least two hours. The pellet was then thawed on ice and resuspended in Bacterial Protein Extraction Reagent (B-PER) (ThermoFisher) with 5 mM MgCl_2_, 1 mM MnCl_2_, 100 µM CaCl_2_, 1 mM (tris(2-carboxyethyl)phosphine) (TCEP) (ThermoFisher), EDTA-free protease inhibitor (Roche), 2 µL/mL DNaseI (Roche), and 2 µL/mL lysozyme (ThermoFisher) for lysis. The cell lysate was then centrifuged at 27,000 × *g* for 30 minutes to separate soluble proteins from insoluble proteins. The soluble cell fraction was decanted, incubated with 20 mM imidazole (ThermoFisher) on ice for 1 hour with nutation, centrifuged at 4000 × *g* for 10 minutes at 4 °C and decanted again. The supernatant was purified by nickel chromatography using an AKTA FPLC (Prime Plus) with HisPur Ni-NTA resin (ThermoFisher) in FPLC buffer (20 mM HEPES, 100 mM KCl, 1 mM TCEP, 1 mM MgCl_2_, 5% glycerol, pH 7.5) with an increasing imidazole gradient up to 400 mM. The elution peak fractions were pooled and applied to a size exclusion chromatography with HiPrep Sephacryl S-100 HR column (Cytiva) column equilibrated with FPLC buffer. Pure fractions were pooled and concentrated with a 10 KDa molecular weight cut-off (MWCO) Amicon Ultra centrifugal filter (Millipore Sigma) to working stock concentrations of 5 µM, 25 µM, and 51 µM after running a denaturing 4-20% SDS-PAGE gel (BioRad) to confirm protein size and purity. Finally, the protein samples were flash frozen in liquid N_2_ and stored at -80 °C.

### X-ray crystallography sample preparation

*E. coli* PFK (10 mg/mL) was incubated at 4 °C for 12 h with 0.1 M sodium citrate. After incubation, the mixture was centrifuged (14,000 rpm, 4 °C, 10 min), and crystals were grown by hanging-drop vapor diffusion using 400 nL of the protein solution and 300 nL of reservoir solution containing 0.1 M Tris pH 8.5, 0.3 M magnesium formate dihydrate (Index HR2-144 solution 16, Hampton Research) at 20 °C.

### X-ray crystallography data collection and model building

Diffraction data were collected at the NYX beamline 19-ID of the National Synchrotron Light Source II (NSLS-II) at Brookhaven National Laboratory. The raw diffraction HDF5 images were processed using XDS to perform indexing, integration and initial scaling.^61^ Data scaling and merging were carried out using the CCP4-supported program, AIMLESS.^62^ Molecular replacement was performed with Phaser, using an AlphaFold3 model as the search model for the structure.^63,64^ Coot was used for manual modelling.^65^ Structural refinement was conducted using both Phenix.refine and REFMAC.^66,67^ Diffraction data and structure refinement statistics are summarized in Table S6.

### HX/MS experiments

#### Sample preparation

H_2_O exchange buffer (50 mM HEPES, 100 mM KCl, 10 mM MgCl_2_, 1 mM TCEP, pH 7.5) was lyophilized in 5 mL aliquots using a Labconco FreeZone 4.5 lyophilizer for at least 24 h, and then re-hydrated in 5 mL D_2_O to produce deuterated exchange buffer. Before each HX/MS experiment, purified EcPFK samples were thawed on ice and buffer-exchanged using a 10 KDa molecular weight cut-off centrifugal filter (Pierce) to HX/MS buffer without glycerol (50 mM HEPES, 100 mM KCl, 10 mM MgCl_2_, 1 mM TCEP, pH 7.5). The sample was then subjected to an additional filtration step with a 0.22 µm spin column (MilliporeSigma) to separate out any precipitates. Stock solutions of EcPFK for apo, ADP, and PEP conditions were prepared to a concentration of 23.5 µM EcPFK, with 96 mM ADP (MilliporeSigma) or PEP (MilliporeSigma), and stored at constant temperature on the LEAP (Trajan) autosampler tray until injection.

#### Preparation of fully deuterated control samples

EcPFK samples were thawed on ice and buffer-exchanged, as described previously, to a non-glycerol buffer (50 mM HEPES, 100 mM KCl, 10 mM MgCl_2_, 1 mM TCEP, pH 7.5) to a concentration of 39 µM. Lyophilized buffer salts and guanidine hydrochloride (GuHCl) (Sigma-Aldrich) were dissolved in D₂O to yield a denaturing buffer containing 3 M GuHCl. The samples were then diluted ten-fold in the denaturing buffer (50 µL protein in 450 µL denaturing buffer) and incubated for 24 hours at 37 °C. The exchanged solution was manually substituted for an equivalent volume of the exchange reaction in the standard HX/MS workflow, using a quench buffer consisting of 3% acetonitrile (Fisher Scientific) and 1% formic acid (Fisher Scientific) without GuHCl to maintain the composition of the quenched exchange reaction.

#### HX/MS data collection

The LEAP system (Trajan) was used for automated HX liquid handling and MS injections with the Chronos 5.8.3 software (Trajan). For each sample at each timepoint, 75 µL of exchange buffer was added to 7.5 µL sample stock solution and mixed by pipetting. After each sample reached its deuteration timepoint, 75 µL of the mix was added to 75 µL of quench solution, consisting of 3 M GuHCl (Sigma-Aldrich) in 3% acetonitrile (Fisher Scientific) and 1% formic acid (Fisher Scientific), chilled to 2 °C and mixed. Subsequently, 140 µL of the chilled, quenched mixture was injected into the protease column (fungal protease type XIII, nepenthesin or alanyl aminopeptidase as indicated in **Table S1**), which was chilled to 7 °C. Samples were allowed to exchange a series of timepoints as indicated in **Table S2**. Eluted peptides entered the mass spectrometer (Bruker Maxis II ETD ESI-QqTOF) via electrospray ionization. For each sample, tandem mass spectrometry (MS/MS) experiments were performed with the same MS settings for peptide identification. Bruker Compass Hystar 5.1 software was used to acquire the data.

#### HX/MS data analysis

HX/MS data from three biological replicates and multiple protease columns (fungal protease type XIII, nepenthesin or alanyl aminopeptidase; Affipro) were analyzed with PIGEON, an HX/MS disambiguation software.^38^ We collected 2-3 MS/MS datasets per HX/MS experiment and produced a compound list in .mgf format with Bruker Compass DataAnalysis software. MS1 and MS2 peaks are matched to theoretical peptide and fragment *m*/*z* values and a score is assigned to each match. The .mgf files were then input into PIGEON to produce pooled peptide sources to use as input for the HDExaminer v3.3 software (Trajan). HDExaminer v3.3 was used to identify the mass spectrum for each time point and peptide in the experiment and then manually checked to ensure the automatically selected retention times matched the same chromatographic peak for each time point. Residue-level free energies of opening (ΔG_op_), associated confidence values, and differences in free energies of opening between states (ΔΔG_op_) were calculated by PFNet.^40^ PFNet outputs a residue-level confidence score on a 0–1 scale, obtained by mapping the predicted absolute error in ΔG_op_ to confidence using a single exponential decay function, such that lower predicted uncertainty corresponds to higher confidence. These energies were mapped onto the crystal structure of ADP-bound EcPFK (PDB: 1PFK) and visualized using PyMOL (Schrödinger), a molecular visualization software. For nonsingle-resolved resolution segments, PFNet outputs exchangeable ΔG_op_ values. These segment-averaged ΔG_op_ values were assigned to each residue within the segment to facilitate PyMOL visualization.

### Activity assay

EcPFK activity was measured using an enzyme-coupled assay adapted from Phong et al., 2013, in which fructose-1,6-bisphosphate formation is coupled to NADH oxidation.^36^ Assays were performed in a total volume of 100 µL in half-area UV-transparent 96-well plates (Grenier). Each well contained a final concentration of 0.3 mM NADH (Millipore Sigma), 1 U/mL aldolase (Millipore Sigma), 1U/rxn glycerophosphate dehydrogenase-triosephosphate isomerase (a-GDH-TPI) (Millipore Sigma), 0.5 mM ATP (Millipore Sigma), and a range of Fructose-6-phosphate (F6P) (ThermoFisher) concentrations, in activity assay buffer (50 mM HEPES, 100 mM KCl, 10 mM MgCl_2_, 1 mM TCEP, 7.5% glycerol, pH 7.5). Auxiliary enzymes (a-GDH-TPI) were de-salted using a centrifugal filter (10 kDa MWCO, Pierce) before use. Assay plates were equilibrated to 30 °C for 10 minutes before reactions were initiated by addition of EcPFK. The final concentration of EcPFK, prepared as previously described, was 10 nM. Final concentrations of ADP (MilliporeSigma) were 0.1 and 0.3 mM, and final concentrations of PEP (MilliporeSigma) were 2 and 5 mM. Final concentrations of F6P ranged from 0.1 to 2.0 mM. NADH oxidation was monitored by measuring absorbance at 340 nm at 30 °C using a Biotek Synergy Neo2 plate reader (Agilent), with readings collected every 30 seconds for 15 minutes. EcPFK activity was quantified as the initial rate of NADH consumption and plotted as a function of F6P concentration in the absence and presence of allosteric ligands. Each condition was measured in three technical replicates. Mean values and standard deviations, represented as error bars in **Fig. 1**, were calculated across replicates. For plotting, initial rates were normalized to the maximal initial velocity within each condition. This quantification was done using the SciPy python library.^68^

### Native gel electrophoresis

Native gel electrophoresis was performed to assess the oligomerization state of purified enzyme under different conditions and in the presence of effectors of interest. Frozen EcPFK stock samples (25 µM) were thawed and prepared as described previously. Ligands in excess concentrations together with sample buffer (20 mM HEPES, 100 mM KCl, 1 mM TCEP, 1 mM MgCl_2_, 5% glycerol, pH 7.5) were diluted to a final volume of 30 µL and a final EcPFK concentration of 2.5 µM. Final ligand concentrations were 100 mM isocitrate (ThermoFisher), 100 mM fumarate (ThermoFisher), 100 mM sodium citrate (Fisher), 20 mM PEP (ThermoFisher), 8 mM ATP (Millipore Sigma), 8 mM F6P (ThermoFisher), and 1.2 mM ADP (Millipore Sigma). Samples were incubated on ice for 15 minutes prior to loading. Following incubation, samples were diluted two-fold in native sample buffer (BioRad) and loaded into a 7.5% stain-free gel (BioRad). NativeMark unstained protein standard (Invitrogen) was loaded in parallel. Electrophoresis was carried out at 90-100 V in Tris-Glycine running buffer. Gels were subsequently stained with Coomassie blue (Pierce) and imaged using a BioRad Chemi Doc XRS+ imager.

### Mass photometry

Mass photometry measurements were performed using a Refeyn OneMP instrument (Refeyn Ltd., Oxford, UK) and data acquisition performed using AcquireMP v2.3.0 software (Refeyn Ltd.). EcPFK stock solutions were thawed and buffer-exchanged into activity assay buffer (20 mM HEPES, 100 mM KCl, 1 mM TCEP, 1 mM MgCl2, pH 7.5) as described previously to remove glycerol from the samples. The sample was then filtered with a 0.2 µm spin filter (MilliporeSigma). For each measurement, microscope coverslips (Refeyn Ltd.) were prepared immediately prior to use and fitted with a silicone template (Millipore Sigma) to form individual reaction chambers. Following instrument calibration, 2 µL protein sample was diluted in 10 µL activity assay buffer to yield a final concentration of 0.24 µM and mixed thoroughly. Mass photometry signals were recorded for 60 s, allowing detection of at least 2 × 10^3^ individual protein molecules per measurement. The data were analyzed using the Refeyn AcquireMP v2.3.0 software.

### Protein structural modeling

X-ray crystal structures were used to model EcPFK in apo and PEP-bound states. Structures were downloaded from the Research Collaboratory for Structural Bioinformatics (RCSB) Protein Data Bank (PDB) in PDB format and relaxed with Rosetta, a macromolecular modeling and design software, with coordinate constraints applied to the starting structure (using the relax application option *-relax:constrain_relax_to_start_coords*).^69–71^ This protocol was applied to the ADP-bound EcPFK (PDB: 1PFK) and to the models prepared for subsequent MD simulations, as described below.

### Molecular dynamics simulations

Input models for MD simulations were derived from the X-ray crystal structure of ADP-bound *E. coli* PFKA (PDB: 1PFK). Because a full-length apo EcPFK crystal structure was not yet available to us, we generated an apo model by removing ligands from the ADP-bound structure followed by a constrained relax as described above. We constructed the PEP-bound model by aligning the structure of the *B. stearothermophilus* PFK-PEP complex (PDB: 4I4I),^23^ the only PEP-bound bacterial structure to our knowledge, onto the apo EcPFK model, and adding the PEP molecule to each E site in the tetrameric structure. The resulting model was then relaxed as described above. The CHARMM36 forcefield was applied and we performed the simulations with the GROMACS package (version 2022).^72–74^ The protein systems were energy-minimized using the steepest descent minimization method until the maximum force was less than 1000 kJ/mol/nm^2^, followed by 100 ps restrained MD simulations in NVT ensemble by constraining the heavy atoms of the protein and ligand at 298.15 K. We further equilibrated the resulting simulation system for a 1000 ps NPT simulation at 1 atm and 298.15 K with 1000 kJ/mol/nm^2^ position restraints on the protein and ligand, followed by softer 100 kJ/mol/nm^2^ position restraints on ligand for 1000 ps NPT simulation at the same condition. Finally, we conducted the NPT production MD simulations in the same conditions without restriction, using a time step of 2 fs, a nonbonded cutoff of 12 Å, and particle-mesh Ewald longrange electrostatics. Three 3 µs simulations were collected for each system as initial datasets. Control simulations of EcPFK with ADP bound in both the E and active sites were also performed. Three replicates of 1 µs simulations were collected as previously described.

### Adaptive sampling

Adaptive sampling was conducted from input structures (seeds) located at the edge of the FEL from initial MSMs (using backbone torsions as the input feature) built from 9 µs MD data (**Fig. S4**) (see next section). Multiple seeds were selected for each system. The protein and ligand complex were extracted, re-solvated in a new simulation box, and simulated at identical conditions for 1000 or 100 ns. This process was repeated until the FEL and implied time scale analysis converged (**Fig. S4**). Finally, 30 µs, 29 µs, 30 µs data were collected for apo EcPFK, EcPFK-ADP and EcPFK-PEP, respectively.

### Markov state models

The MSMs were built from MD trajectories using PyEMMA (2.5.12).^75^ Secondary structure features were selected as the input features (**Fig. S6**). We used time-lagged independent component analysis (TICA) to project the original high dimensional data to a subspace that captures the slow motions of the protein. Then, we applied the K-means algorithm to cluster tICA data to discrete trajectories of the microstates. We determined the number of K-means clusters as the lowest number of clusters for achieving a converged VAMP2 core analysis.^42^ Then, we used the discrete trajectory for each functional state of EcPFK to estimate the MSM for that state using a Bayesian framework implemented in PyEMMA.^76^ The lag time τ was chosen based on the convergence of implied timescales for each system. Thus, the final models were built using 100 microstates, a lag time of 30 ns for apo EcPFK, EcPFK-ADP and EcPFK-PEP systems (**Fig. S7)**. Then, we applied the PCCA+ method implemented in PyEMMA to lump microstates with faster inner transitions into larger metastable states.^77,78^ The model was validated by the Chapman-Kolmogorov test by computing the transition probability between metastable states for different lagtimes.^44^ The results show that the predictions of the final model at kτ are within the statistical error of the independently estimated model at kτ (**Fig. S7**).

### hPFK simulations

hPFK simulations were performed on Anton 2.^51^ System preparation and pre-equilibration followed a procedure similar to that described in the molecular dynamics simulations section above using GROMACS with the Amber ff14SB force field. The initial models were based on X-ray crystal structures of hPFK (PDB IDs: 4XYJ and 4XYK), which were subjected to 100 ns of equilibration on GPU hardware. The equilibrated GROMACS systems were subsequently converted to Anton-compatible formats. Topology and coordinate files were transformed using InterMol,^79^ followed by conversion to DMS format and force field assignment using viparr tool by D.E. Shaw Research with the Amber ff14SB^80^ parameters and TIP3P water model. The resulting DMS files were used as inputs for Anton 2 production simulations. For each system, a 4 µs production trajectory was collected.

### Pocket volume analysis

MDpocket was used to detect small-molecule binding sites and calculate pocket volumes on the MSM-reweighted ensembles for apo, ADP-bound, and PEP-bound EcPFK.^81^ Prior to analysis, all MD frames were aligned to the EcPFK X-ray crystal structure (PDB: 1PFK). Active site pockets were manually selected for each ensemble using frequency maps at an isovalue of 0.7, corresponding to at least 70% of pocket opening in all frames. Probe spheres within 3 Å of FBP in the active site, based on the aligned crystal structure coordinates, were selected to calculate pocket volume. PyMOL was used to visualize MDpocket-derived probe spheres.

### RMSF analysis

We used MDAnalysis, an open-source tool for analyzing MD simulation data, to calculate residue-level heavy-atom RSMF for each reweighted ensemble. For each trajectory, backbone atom coordinates in every frame were aligned to a reference structure, the crystal structure of ADP-bound EcPFK (PDB: 1PFK), which is also the same starting structure for all simulations in each state. We then computed heavy-atom RMSF values for each residue over the trajectories and averaged over the heavy atoms within each residue. Differences in residue-level RMSF between ensembles (ΔRMSF_APO-HOLO_) were calculated by first averaging values across chains and computing 95% confidence intervals using the Pingouin package.^82^ These differences were visualized by mapping the values onto the crystal structure of ADP-bound EcPFK (PDB: 1PFK). We assessed statistical significance by computing *p*-values using Welch’s unpaired *t*-test using the SciPy python library.^68^

For RMSF analysis of each hPFK lobe, the full-length hPFK amino acid sequence was partitioned into an N- and C-terminal lobe based on inspection of the hPFK crystal structure (PDB: 4XYJ) compared to the EcPFK monomer (PDB: 1PFK). Sequence and structural alignment between lobes was performed using US-align.^52^ We calculated mean RMSF values by averaging RMSF across chains for each residue, calculated the corresponding 95% confidence intervals using Pingouin, and used the SciPy python library to compute Pearson correlation coefficients and to assess whether differences in average RMSF between lobes was significant. Per-residue RSMF was visualized and plotted according to the lobe-lobe sequence alignment. Using the same approach, sequence alignments between EcPFK and each hPFK lobe were generated and used to compute correlations between aligned residues between aligned residues. These correlations were evaluated both globally and within specific key functional sites (as defined in **Table S4**) to assess conservation of dynamics between EcPFK and hPFK.

### RMSD analysis

We calculated per-residue RMSD for each reweighted ensemble using the molecular dynamics analysis software MDTraj.^83^ For each frame, backbone atoms were aligned to a reference structure. We then computed heavy-atom RMSD values for each frame and each residue relative to this reference and visualized per-residue RMSD distributions using python.

To analyze secondary structure RMSD, residues corresponding to each secondary structure element were first defined using the Dictionary of Secondary Structure in Proteins (DSSP) algorithm^42^ applied to the EcPFK crystal structure (PDB: 1PFK, **Table S3**). Secondary structure features were then extracted from each MSM using PyEmma. Distributions of secondary structure RMSD were visualized using python, and Kullback-Leibler (KL) divergence was calculated with the SciPy python library to identify secondary structure elements with large differences in RSMD distribution between ensembles.^68^

### Interaction network analysis

We performed hydrogen bond analysis on each reweighted ensemble using MDAnalysis.^84^ Hydrogen bonds were defined based on geometric criteria, with a donor–acceptor distance ≤ 3.5 Å and a donor-hydrogen-acceptor angle ≥ 150°. For each identified hydrogen bond, the occupancy was computed over the trajectories of the reweighted ensembles. Standard deviations were calculated across the four protomers. For each state-to-state comparison, we retained hydrogen bonds only if the absolute difference in mean occupancy exceeded 10% and was greater than the corresponding standard deviation.

Nonpolar contact and salt bridge analyses were conducted similarly using the following geometric criteria. Nonpolar contacts were defined by a closest heavy-atom distance ≤ 4.5 Å between side-chain atoms of ALA, VAL, LEU, ILE, MET, PHE, and TRP residues, excluding backbone atoms. Salt bridges were defined by a closest-atom distance ≤ 4.5 Å between positively charged nitrogen atoms of ARG, LYS, or HIS residues and negatively charged oxygen atoms of ASP or GLU residues. Differences in interaction occupancy between states for each interaction type were visualized using PyMOL.

### Quantification and statistical analysis

Statistical analyses and quantification procedures are reported in the relevant Methods sections. For HX/MS experiments, residue-level free energies of opening (ΔG_op_) and differences between states (ΔΔG_op_), and associated confidence values were calculated using PFNet, as described above.^40^ EcPFK activity assays were quantified as the initial rate of NADH consumption, averaged across three technical replicates (mean ± SD), normalized to the maximal velocity with each condition, via the SciPy Python library. For MD simulations, ensemble-averaged quantities, including RSMF, RMSD, interaction occupancies, pocket volumes, and secondary structure metrics, were computed from MSM-reweighted trajectories, except for the hPFK simulations for which these quantities were computed directly from raw trajectories. Where applicable, these values were averaged across protomers, and variability is reported as standard deviation or 95% confidence intervals with Pingouin, as indicated. We assessed statistical significance using Welch’s t-tests and performed correlation analyses using Pearson correlation coefficients in Python using SciPy and Pingouin, unless otherwise specified. Significance thresholds and exact values of sampled size (*n*) are reported in the corresponding figure legends.

### Supporting information

Supporting figures and tables are available in the Supplementary PDF (Figures S1-S14, Tables S1-S6, and supplemental references). Supplemental datasets can be found online at <u>Zenodo</u> (DOI 10.5281/zenodo.18390796).

## REFERENCES

1. Berg, J.M., Tymoczko, J.L., and Stryer, L. (2002). Biochemistry, Fifth Edition (W.H. Freeman).

2. Poorman, R.A., Randolph, A., Kemp, R.G., and Heinrikson, R.L. (1984). Evolution of phosphofructokinase—gene duplication and creation of new effector sites. Nature 309, 467–469.

3. Blangy, D. (1968). Phosphofructokinase from *E. Coli* : Evidence for a tetrameric structure of the enzyme. FEBS Lett. 2, 109–111.

4. Webb, B.A., Forouhar, F., Szu, F.-E., Seetharaman, J., Tong, L., and Barber, D.L. (2015). Structures of human phosphofructokinase-1 and atomic basis of cancer-associated mutations. Nature 523, 111–114.

5. Nakajima, H., Raben, N., Hamaguchi, T., and Yamasaki, T. Phosphofructokinase Deficiency Past, Present and Future. http://www.eurekaselect.com.

6. Fernandes, P.M., Kinkead, J., McNae, I., Michels, P.A.M., and Walkinshaw, M.D. (2020). Biochemical and transcript level differences between the three human phosphofructokinases show optimisation of each isoform for specific metabolic niches. Biochem. J. 477, 4425–4441.

7. Frieden, C., Gilbert, H.R., and Bock, P.E. (1976). Phosphofructokinase. III. Correlation of the regulatory kinetic and molecular properties of the rabbit muscle enzyme. J. Biol. Chem. 251, 5644–5647.

8. Bock, P.E., and Frieden, C. (1976). Phosphofructokinase. I. Mechanism of the pH-dependent inactivation and reactivation of the rabbit muscle enzyme. J. Biol. Chem. 251, 5630–5636.

9. Bock, P.E., and Frieden, C. (1976). Phosphofructokinase. II. Role of ligands in pH-dependent structural changes of the rabbit muscle enzyme. J. Biol. Chem. 251, 5637–5643.

10. Schöneberg, T., Kloos, M., Brüser, A., Kirchberger, J., and Sträter, N. (2013). Structure and allosteric regulation of eukaryotic 6-phosphofructokinases. bchm 394, 977–993.

11. Monod, J., Wyman, J., and Changeux, J.-P. (1965). On the nature of allosteric transitions: A plausible model. J. Mol. Biol. 12, 88–118.

12. Tat-Kwong Lau, F., and Fersht, A.R. (1987). Conversion of allosteric inhibition to activation in phosphofructokinase by protein engineering. Nature 326, 811–812.

13. Lau, F.T.K., and Fersht, A.R. (1989). Dissection of the effector-binding site and complementation studies of Escherichia coli phosphofructokinase using site-directed mutagenesis. Biochemistry 28, 6841–6847.

14. Rypniewski, W.R., and Evans, P.R. (1989). Crystal structure of unliganded phosphofructokinase from Escherichia coli. J. Mol. Biol. 207, 805–821.

15. Shirakihara, Y., and Evans, P.R. (1988). Crystal structure of the complex of phosphofructokinase from Escherichia coli with its reaction products. J. Mol. Biol. 204, 973–994.

16. Blangy, D., Buc, H., and Monod, J. (1968). Kinetics of the allosteric interactions of phosphofructokinase from Escherichia coli. J. Mol. Biol. 31, 13–35.

17. Daily, M.D., and Gray, J.J. (2009). Allosteric Communication Occurs via Networks of Tertiary and Quaternary Motions in Proteins. PLOS Comput. Biol. 5, e1000293.

18. Ghode, A., Gross, L.Z., Tee, W.-V., Guarnera, E., Berezovsky, I.N., Biondi, R.M., and Anand, G.S. (2020). Synergistic allostery in multiligand-protein interactions. Biophys. J. 119, 1833–1848.

19. Evans, P.R., and Hudson, P.J. (1979). Structure and control of phosphofructokinase from Bacillus stearothermophilus. Nature 279, 500–504.

20. Evans, P.R., Farrants, G.W., and Hudson, P.J. (1981). Phosphofructokinase: structure and control. Philos. Trans. R. Soc. Lond. B Biol. Sci. 293, 53–62.

21. Schirmer, T., and Evans, P.R. (1990). Structural basis of the allosteric behaviour of phosphofructokinase. Nature 343, 140–145.

22. Kimmel, J.L., and Reinhart, G.D. (2000). Reevaluation of the accepted allosteric mechanism of phosphofructokinase from Bacillus stearothermophilus. Proc. Natl. Acad. Sci. 97, 3844–3849.

23. Mosser, R., Reddy, M.C.M., Bruning, J.B., Sacchettini, J.C., and Reinhart, G.D. (2013). Redefining the Role of the Quaternary Shift in Bacillus stearothermophilus Phosphofructokinase. Biochemistry 52, 5421–5429.

24. Whitaker, A.M., Naik, M.T., Mosser, R.E., and Reinhart, G.D. (2019). Propagation of the Allosteric Signal in Phosphofructokinase from *Bacillus stearothermophilus* Examined by Methyl-Transverse Relaxation-Optimized Spectroscopy Nuclear Magnetic Resonance. Biochemistry 58, 5294–5304.

25. Deville-Bonne, D., and Garel, J.R. (1992). A conformational transition involved in antagonistic substrate binding to the allosteric phosphofructokinase from Escherichia coli. Biochemistry 31, 1695–1700.

26. Johnson, J.L., and Reinhart, G.D. (1997). Failure of a Two-State Model To Describe the Influence of Phospho (*enol*) pyruvate on Phosphofructokinase from *Escherichia coli*. Biochemistry 36, 12814–12822.

27. Pham, A.S., and Reinhart, G.D. (2003). Quantification of Allosteric Influence of Escherichia coli Phosphofructokinase by Frequency Domain Fluorescence. Biophys. J. 85, 656–666.

28. Fenton, A.W., and Reinhart, G.D. (2009). Disentangling the Web of Allosteric Communication in a Homotetramer: Heterotropic Inhibition in Phosphofructokinase from *Escherichia coli*. Biochemistry 48, 12323–12328.

29. Glasgow, A., Hobbs, H.T., Perry, Z.R., Wells, M.L., Marqusee, S., and Kortemme, T. (2023). Ligand-specific changes in conformational flexibility mediate long-range allostery in the lac repressor. Nat. Commun. 14, 1179.

30. Motlagh, H.N., Wrabl, J.O., Li, J., and Hilser, V.J. (2014). The ensemble nature of allostery. Nature 508, 331–339.

31. Henzler-Wildman, K., and Kern, D. (2007). Dynamic personalities of proteins. Nature 450, 964–972.

32. Le Bras, G., and Garel, J.R. (1982). A proteolyzed derivative of E. coli phosphofructokinase is no longer sensitive to allosteric effectors and still shows cooperativity in substrate binding. Biochemistry 21, 6656–6660.

33. Tian, T., Wang, C., Wu, M., Zhang, X., and Zang, J. (2018). Structural Insights into the Regulation of *Staphylococcus aureus* Phosphofructokinase by Tetramer–Dimer Conversion. Biochemistry 57, 4252–4262.

34. McGresham, M.S., Lovingshimer, M., and Reinhart, G.D. (2014). Allosteric Regulation in Phosphofructokinase from the Extreme Thermophile *Thermus thermophilus*. Biochemistry 53, 270–278.

35. Berger, S.A., and Evans, P.R. (1992). Site-directed mutagenesis identifies catalytic residues in the active site of Escherichia coli phosphofructokinase. Biochemistry 31, 9237–9242.

36. Phong, W.Y., Lin, W., Rao, S.P.S., Dick, T., Alonso, S., and Pethe, K. (2013). Characterization of Phosphofructokinase Activity in Mycobacterium tuberculosis Reveals That a Functional Glycolytic Carbon Flow Is Necessary to Limit the Accumulation of Toxic Metabolic Intermediates under Hypoxia. PLoS ONE 8.

37. Konermann, L., Pan, J., and Liu, Y.-H. (2011). Hydrogen exchange mass spectrometry for studying protein structure and dynamics. Chem. Soc. Rev. 40, 1224–1234.

38. Lu, C., Wells, M.L., Reckers, A., McBride, S.K., and Glasgow, A. (2026). Site-resolved energetic information from HX–MS experiments. Nat. Chem. Biol. 22, 307–317.

39. Wells, M.L., Lu, C., Sultanov, D., Weber, K.C., Ahern, E., Gong, Z., Chen, E., and Glasgow, A. (2025). Distinct energetic blueprints diversify function of conserved protein folds. Accepted, Nat. Chem., BioRxiv 2025.04.02.646877.

40. Lu, C., Weber, K.C., McBride, S.K., Reckers, A., and Glasgow, A. A machine learning method for calculating highly localized protein stabilities. Under review, BioRxiv 2025.10.21.683809.

41. Rodriguez, D.C.P., Weber, K.C., Sundberg, B., and Glasgow, A. (2024). MAGPIE: An interactive tool for visualizing and analyzing protein–ligand interactions. Protein Sci. 33, e5027.

42. Kabsch, W., and Sander, C. (1983). Dictionary of protein secondary structure: Pattern recognition of hydrogen-bonded and geometrical features. Biopolymers 22, 2577–2637.

43. Wang, W., Cao, S., Zhu, L., and Huang, X. (2018). Constructing Markov State Models to elucidate the functional conformational changes of complex biomolecules. WIREs Comput. Mol. Sci. 8, e1343.

44. Prinz, J.-H., Wu, H., Sarich, M., Keller, B., Senne, M., Held, M., Chodera, J.D., Schütte, C., and Noé, F. (2011). Markov models of molecular kinetics: Generation and validation. J. Chem. Phys. 134.

45. Chodera, J.D., and Noé, F. (2014). Markov state models of biomolecular conformational dynamics. Curr. Opin. Struct. Biol. 25, 135–144.

46. Mardt, A., Pasquali, L., Wu, H., and Noé, F. (2018). VAMPnets for deep learning of molecular kinetics. Nat. Commun. 9, 5.

47. Endres, D.M., and Schindelin, J.E. (2003). A new metric for probability distributions. IEEE Trans. Inf. Theory 49, 1858–1860.

48. Kemp, R.G., and Gunasekera, D. (2002). Evolution of the Allosteric Ligand Sites of Mammalian Phosphofructo-1-kinase. Biochemistry 41, 9426–9430.

49. Martínez-Costa, O.H., Hermida, C., Sánchez-Martínez, C., Santamaría, B., and Aragón, J.J. (2004). Identification of C-terminal motifs responsible for transmission of inhibition by ATP of mammalian phosphofructokinase, and their contribution to other allosteric effects. Biochem. J. 377, 77–84.

50. Kloos, M., Brüser, A., Kirchberger, J., Schöneberg, T., and Sträter, N. (2014). Crystallization and preliminary crystallographic analysis of human muscle phosphofructokinase, the main regulator of glycolysis. Acta Crystallogr. Sect. F Struct. Biol. Commun. 70, 578–582.

51. Shaw, D.E., Grossman, J.P., Bank, J.A., Batson, B., Butts, J.A., Chao, J.C., Deneroff, M.M., Dror, R.O., Even, A., Fenton, C.H., et al. (2014). Anton 2: Raising the Bar for Performance and Programmability in a Special-Purpose Molecular Dynamics Supercomputer. In SC ’14: Proceedings of the International Conference for High Performance Computing, Networking, Storage and Analysis, pp. 41–53.

52. Zhang, C., Shine, M., Pyle, A.M., and Zhang, Y. (2022). US-align: universal structure alignments of proteins, nucleic acids, and macromolecular complexes. Nat. Methods 19, 1109–1115.

53. Pham, A.S., Janiak-Spens, F., and Reinhart, G.D. (2001). Persistent Binding of MgADP to the E187A Mutant of *Escherichia coli* Phosphofructokinase in the Absence of Allosteric Effects. Biochemistry 40, 4140–4149.

54. Fenton, A.W., Paricharttanakul, N.M., and Reinhart, G.D. (2003). Identification of Substrate Contact Residues Important for the Allosteric Regulation of Phosphofructokinase from *Eschericia coli*. Biochemistry 42, 6453–6459.

55. Tarui, S., Ikura, Y., Tanaka, T., Suda, M., and Nishikawa, M. (1965). Phosphofructokinase deficiency in skeletal muscle. A new type of glycogenosis. Biochem. Biophys. Res. Commun. 19, 517–523.

56. Yi, W., Clark, P.M., Mason, D.E., Keenan, M.C., Hill, C., Goddard, W.A., Peters, E.C., Driggers, E.M., and Hsieh-Wilson, L.C. (2012). Phosphofructokinase 1 Glycosylation Regulates Cell Growth and Metabolism. Science 337, 975–980.

57. Moreno-Sánchez, R., Marín-Hernández, A., Gallardo-Pérez, J.C., Quezada, H., Encalada, R., Rodríguez-Enríquez, S., and Saavedra, E. (2012). Phosphofructokinase type 1 kinetics, isoform expression, and gene polymorphisms in cancer cells. J. Cell. Biochem. 113, 1692–1703.

58. Weber, K.C., Lu, C., Alvarez, R.V., Pascal, B.D., and Glasgow, A. (2025). HXMS: a standardized file format for HX/MS data. Under review, BioRxiv 2025.10.14.682397.

59. Andersen, K.R., Leksa, N.C., and Schwartz, T.U. (2013). Optimized *E. coli* expression strain LOBSTR eliminates common contaminants from His-tag purification. Proteins Struct. Funct. Bioinforma. 81, 1857–1861.

60. Engler, C., Kandzia, R., and Marillonnet, S. (2008). A one pot, one step, precision cloning method with high throughput capability. PloS One 3, e3647.

61. Kabsch, W. (2010). XDS. Acta Crystallogr. D Biol. Crystallogr. 66, 125–132.

62. Agirre, J., Atanasova, M., Bagdonas, H., Ballard, C.B., Baslé, A., Beilsten-Edmands, J., Borges, R.J., Brown, D.G., Burgos-Mármol, J.J., Berrisford, J.M., et al. (2023). The CCP4 suite: integrative software for macromolecular crystallography. Acta Crystallogr. Sect. Struct. Biol. 79, 449–461.

63. Abramson, J., Adler, J., Dunger, J., Evans, R., Green, T., Pritzel, A., Ronneberger, O., Willmore, L., Ballard, A.J., Bambrick, J., et al. (2024). Accurate structure prediction of biomolecular interactions with AlphaFold 3. Nature, 1–3.

64. McCoy, A.J., Grosse-Kunstleve, R.W., Adams, P.D., Winn, M.D., Storoni, L.C., and Read, R.J. (2007). Phaser crystallographic software. J. Appl. Crystallogr. 40, 658–674.

65. Emsley, P., and Cowtan, K. (2004). Coot: model-building tools for molecular graphics. Acta Crystallogr. D Biol. Crystallogr. 60, 2126–2132.

66. Murshudov, G.N., Skubák, P., Lebedev, A.A., Pannu, N.S., Steiner, R.A., Nicholls, R.A., Winn, M.D., Long, F., and Vagin, A.A. (2011). REFMAC5 for the refinement of macromolecular crystal structures. Biol. Crystallogr. 67, 355–367.

67. Afonine, P.V., Grosse-Kunstleve, R.W., Echols, N., Headd, J.J., Moriarty, N.W., Mustyakimov, M., Terwilliger, T.C., Urzhumtsev, A., Zwart, P.H., and Adams, P.D. (2012). Towards automated crystallographic structure refinement with phenix. refine. Biol. Crystallogr. 68, 352–367.

68. Virtanen, P., Gommers, R., Oliphant, T.E., Haberland, M., Reddy, T., Cournapeau, D., Burovski, E., Peterson, P., Weckesser, W., and Bright, J. (2020). SciPy 1.0: fundamental algorithms for scientific computing in Python. Nat. Methods 17, 261–272.

69. Berman, H.M., Westbrook, J., Feng, Z., Gilliland, G., Bhat, T.N., Weissig, H., Shindyalov, I.N., and Bourne, P.E. (2000). The Protein Data Bank. Nucleic Acids Res. 28, 235–242.

70. Conway, P., Tyka, M.D., DiMaio, F., Konerding, D.E., and Baker, D. (2014). Relaxation of backbone bond geometry improves protein energy landscape modeling. Protein Sci. Publ. Protein Soc. 23, 47–55.

71. Leaver-Fay, A., Tyka, M., Lewis, S.M., Lange, O.F., Thompson, J., Jacak, R., Kaufman, K.W., Renfrew, P.D., Smith, C.A., and Sheffler, W. (2011). ROSETTA3: an object-oriented software suite for the simulation and design of macromolecules. In Methods in enzymology (Elsevier), pp. 545–574.

72. Abraham, M.J., Murtola, T., Schulz, R., Páll, S., Smith, J.C., Hess, B., and Lindahl, E. (2015). GROMACS: High performance molecular simulations through multi-level parallelism from laptops to supercomputers. SoftwareX 1–2, 19–25.

73. Vanommeslaeghe, K., Hatcher, E., Acharya, C., Kundu, S., Zhong, S., Shim, J., Darian, E., Guvench, O., Lopes, P., Vorobyov, I., et al. (2010). CHARMM General Force Field (CGenFF): A force field for drug-like molecules compatible with the CHARMM all-atom additive biological force fields. J. Comput. Chem. 31, 671–690.

74. Huang, J., and MacKerell, A.D. (2013). CHARMM36 all-atom additive protein force field: Validation based on comparison to NMR data. J. Comput. Chem. 34, 2135–2145.

75. Scherer, M.K., Trendelkamp-Schroer, B., Paul, F., Pérez-Hernández, G., Hoffmann, M., Plattner, N., Wehmeyer, C., Prinz, J.-H., and Noé, F. (2015). PyEMMA 2: A Software Package for Estimation, Validation, and Analysis of Markov Models. J. Chem. Theory Comput. 11, 5525–5542.

76. Trendelkamp-Schroer, B., Wu, H., Paul, F., and Noé, F. (2015). Estimation and uncertainty of reversible Markov models. J. Chem. Phys. 143, 174101.

77. Röblitz, S., and Weber, M. (2013). Fuzzy spectral clustering by PCCA+: application to Markov state models and data classification. Adv. Data Anal. Classif. 7, 147–179.

78. Noé, F., Wu, H., Prinz, J.-H., and Plattner, N. (2013). Projected and hidden Markov models for calculating kinetics and metastable states of complex molecules. J. Chem. Phys. 139.

79. Shirts, M.R., Klein, C., Swails, J.M., Yin, J., Gilson, M.K., Mobley, D.L., Case, D.A., and Zhong, E.D. (2017). Lessons learned from comparing molecular dynamics engines on the SAMPL5 dataset. J. Comput. Aided Mol. Des. 31, 147–161.

80. Maier, J.A., Martinez, C., Kasavajhala, K., Wickstrom, L., Hauser, K.E., and Simmerling, C. (2015). ff14SB: Improving the Accuracy of Protein Side Chain and Backbone Parameters from ff99SB. J. Chem. Theory Comput. 11, 3696–3713.

81. Schmidtke, P., Bidon-Chanal, A., Luque, F.J., and Barril, X. (2011). MDpocket: open-source cavity detection and characterization on molecular dynamics trajectories. Bioinformatics 27, 3276–3285.

82. Vallat, R. (2018). Pingouin: statistics in Python. J Open Source Softw 3, 1026.

83. McGibbon, R.T., Beauchamp, K.A., Harrigan, M.P., Klein, C., Swails, J.M., Hernández, C.X., Schwantes, C.R., Wang, L.-P., Lane, T.J., and Pande, V.S. (2015). MDTraj: a modern open library for the analysis of molecular dynamics trajectories. Biophys. J. 109, 1528–1532.

84. Michaud-Agrawal, N., Denning, E.J., Woolf, T.B., and Beckstein, O. (2011). MDAnalysis: A toolkit for the analysis of molecular dynamics simulations. J. Comput. Chem. 32, 2319–2327.

