## Supplemental information for "Bidirectional allosteric ligand regulation in a central glycolytic enzyme"

**Table of contents**

**Supplemental figures** 3

Figure S1. R- and T-states in BsPFK vs. EcPFK. 3

Figure S2. Oligomeric states of EcPFK, related to Figure 1. 4

Figure S3. HX/MS analysis of EcPFK functional states, related to Figure 2. 5

Figure S4. MD/MSM construction, related to Figure 3. 7

Figure S5. MSM feature selection metrics for three features, related to Figure 3. 8

Figure S6. Comparing feature robustness of reweighted ensembles, related to Figure 3. 9

Figure S7. MSM construction and validation for EcPFK, related to Figure 3. 11

Figure S8. Comparing MSMs with HX/MS-derived $\Delta G_{op}$, related to Figure 3. 12

Figure S9. RMSF analysis for EcPFK functional states, related to Figure 3. 13

Figure S10. Changes in interactions in reweighted ensembles, related to Figure 4. 15

Figure S11. F188 as a molecular switch, related to Figure 4. 16

Figure S12. Substrate binding in the PEP state, related to Figure 5. 17

Figure S13. Average active site volume for EcPFK functional states, related to Figure 5. 18

Figure S14. Evolutionary origins of hPFK ligand regulation, related to Figure 6. 20

**Supplemental tables** 22

Table S1. HX experiment summary table. 22

Table S2. PFNet results summary. 23

Table S3. DSSP-defined secondary structure segment indices. 24

Table S4. Site definitions key functional areas in PFK, related to Figure 6. 26

Table S5. DNA sequence for plasmid used in this study. 27

Table S6. X-ray crystallography summary statistics. 29

**Supplemental references** 30

**Supplemental figures**

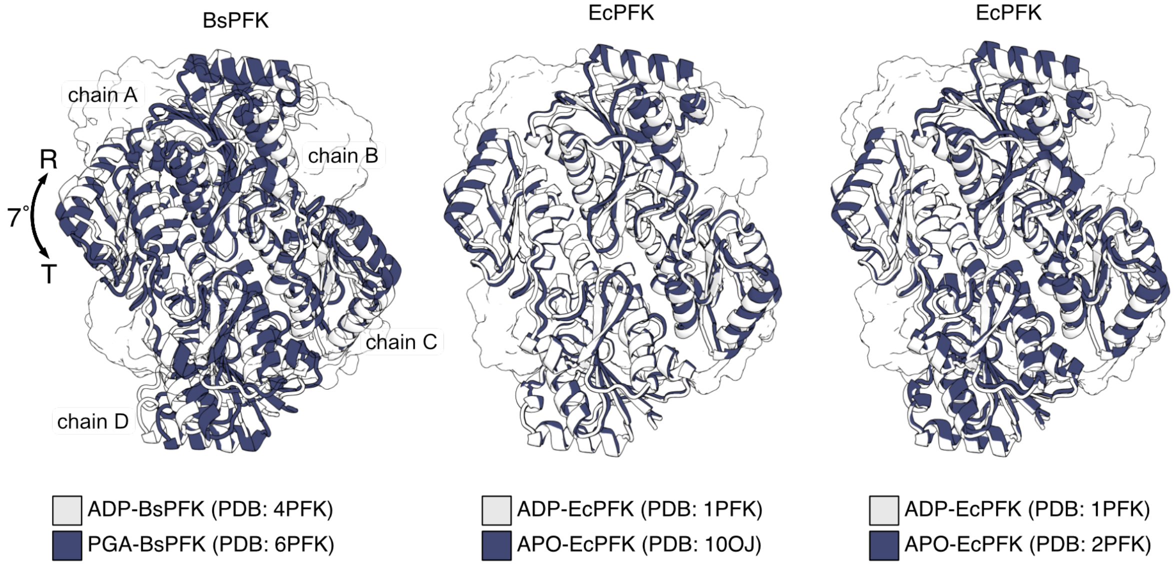


**Figure S1. R- and T-states in BsPFK vs. EcPFK.**

Adapted from Evans and Hudson, 1979, and Schirmer and Evans, 1990 [1, 2]. Previous studies identified a quaternary shift between the ADP-bound structure in the R-state and the PGA-bound structure in the T-state of BsPFK, corresponding to a 7° rotation of one dimer relative to the other. This conformational difference was proposed to be central to the mechanism of BsPFK allosteric regulation and results in the closing of the active site in the T-state. In contrast, when the crystal structure of unliganded EcPFK (expected to resemble the BsPFK T-state conformation) was solved, authors noted that it was more similar to the activator-bound EcPFK structure than the T-state BsPFK structure: "*Compared to the active conformation, the unliganded structure does not show the substantial quaternary structure change seen in the BsPFK T-state structure*," from (Rypniewski and Evans, 1989) [3]. To demonstrate this, shown here are comparisons between the R- and T-state structures of BsPFK and EcPFK (our newly solved apo structure, 10OJ, and the original apo structure, 2PFK): one dimer (chains A and B) in the R state is aligned to the corresponding dimer (chains A and B) in the T state. When the full tetramer (chains A-D) is shown, the 7˚ rotation between dimers is evident in BsPFK, but absent in EcPFK.

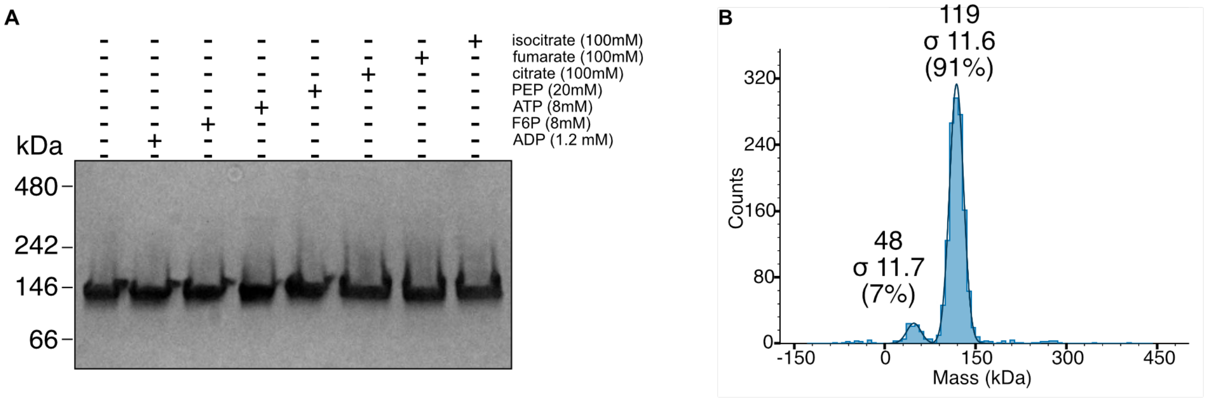


**Figure S2. Oligomeric states of EcPFK, related to Figure 1.**

**(A)** In agreement with previous biochemical studies, native PAGE shows that purified EcPFK does not undergo a change in oligomerization state in response to binding F6P (substrate), ADP (activator), PEP (inhibitor), citrate (inhibitor of eukaryotic PFK), fumarate, or isocitrate.
**(B)** Mass photometry shows that apo EcPFK is primarily a tetramer in solution at 0.24 μM.

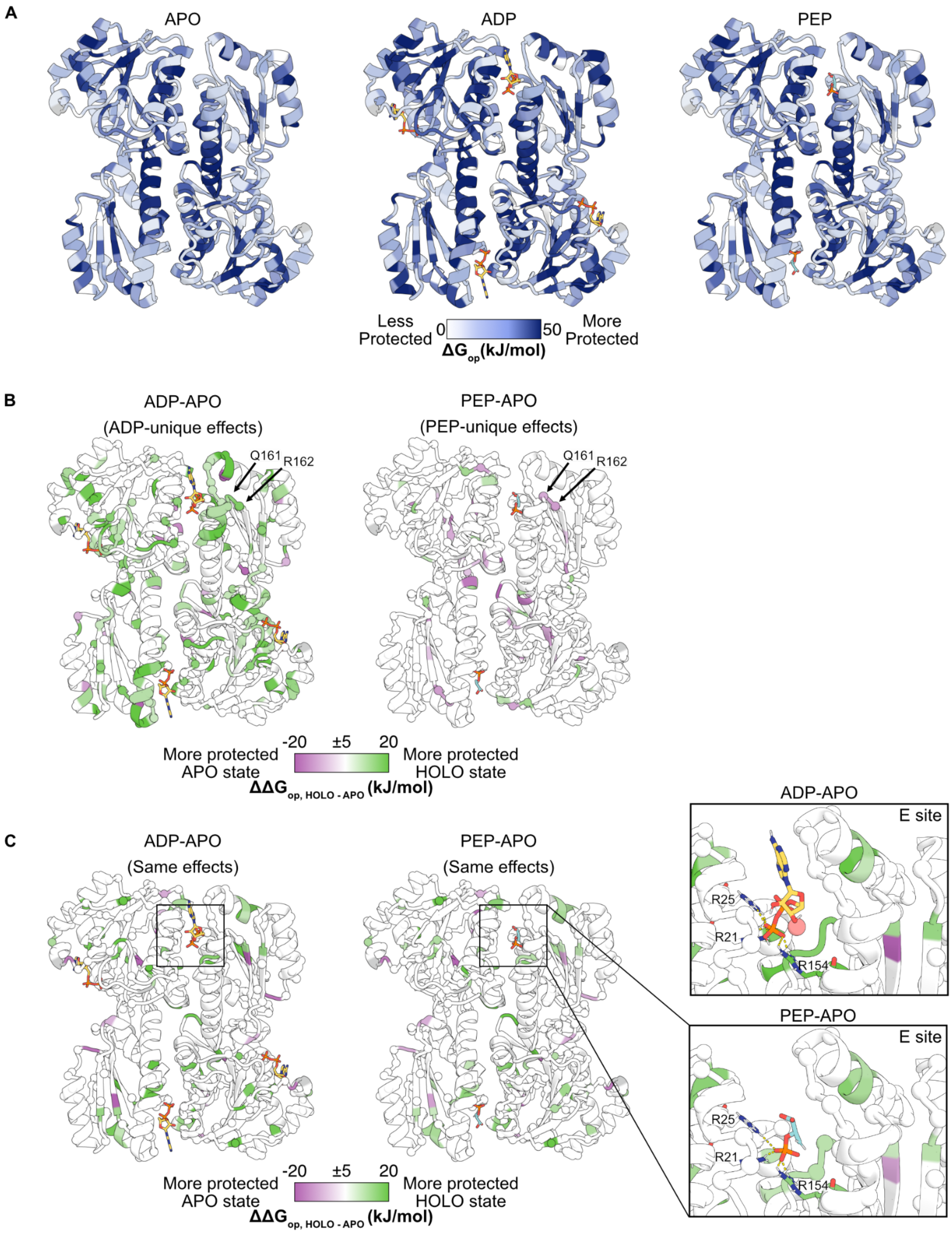


**Figure S3. HX/MS analysis of EcPFK functional states, related to Figure 2.**

**(A)** Residue-level $\Delta G_{op}$ mapped onto the EcPFK crystal structures (PDB: 1PFK).

**(B)** $\Delta\Delta G_{op.\ HOLO\text{-}APO}$ highlighting only ligand-specific energetic effects, *i.e.*, residues that are stabilized in the ADP-bound ensemble relative to the apo state but not stabilized in the PEP-bound ensemble vs. the apo state. Residues showing the opposite responses are also included, *i.e.*, residues stabilized by ADP but destabilized by PEP relative to the apo state. Spheres denote high-confidence residues (PFNet confidence >0.8).
**(C)** Residue-level energetic effects common to both ADP- and PEP-bound ensembles vs. the apo state ensemble.

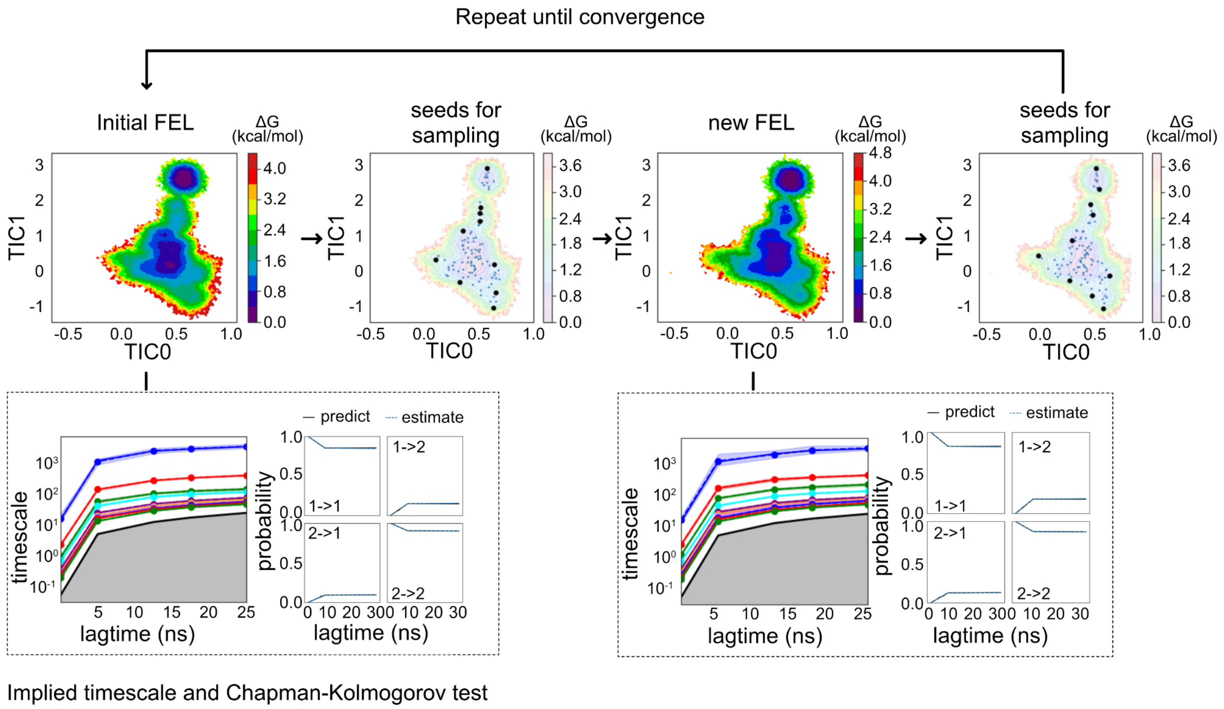


**Figure S4. MD/MSM construction, related to Figure 3.**

Adaptive sampling strategy used in this study, illustrated using the EcPFK-ADP functional state as an example (see Methods).

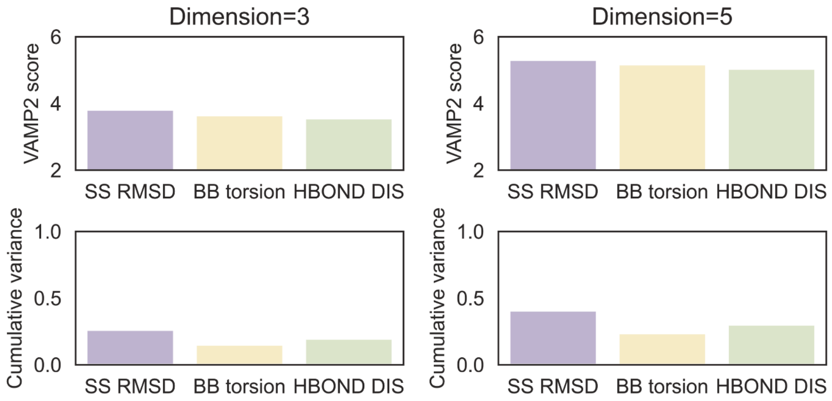


**Figure S5. MSM feature selection metrics for three features, related to Figure 3.**

Feature selection based on VAMP2 scores and cumulative variance (Prinz *et al.*, 2011) [4]. Comparisons across secondary structure segments RMSD (SS RMSD), backbone torsion (BB torsion), and backbone hydrogen bond distance (HBOND DIS) features indicate that SS RMSD consistently achieves the highest VAMP2 scores and explained variance across embedding dimensions. The SS RMSD was therefore selected for subsequent analysis.

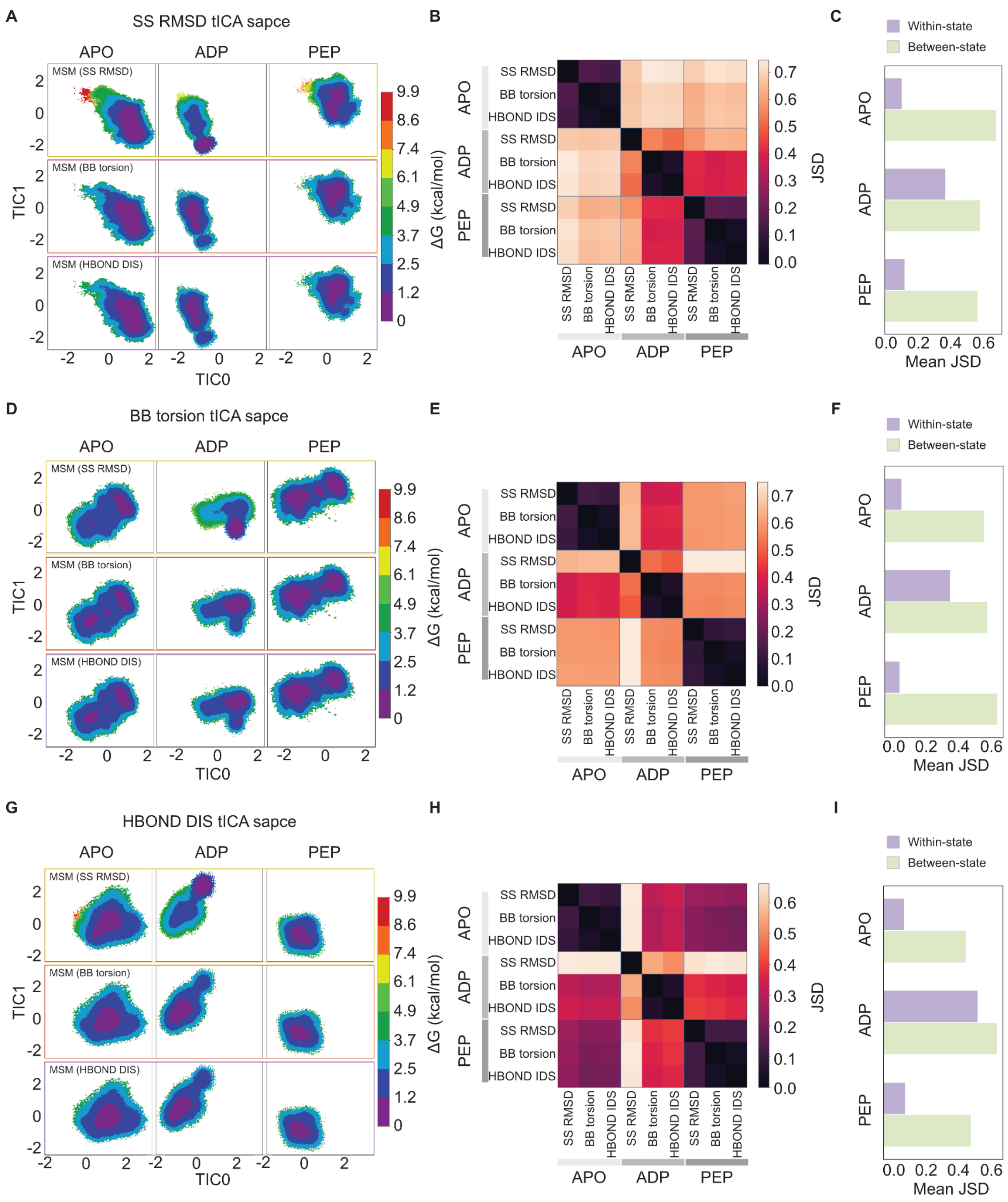


**Figure S6. Comparing feature robustness of reweighted ensembles, related to Figure 3.**

**(A)** MSM-reweighted ensembles projected onto a TICA space (TIC0–TIC1) obtained from the secondary structure segment RMSD (SS RMSD) feature. Rows show MSMs constructed from distinct features including SS RMSD, backbone torsion (BB torsion) and backbone hydrogen bond distance (HBOND DIS) and columns corresponding to the APO, ADP, and PEP states

**(B)** Jensen-Shannon divergence (JSD) (Endres *et al.*, 2003) [5] between MSM-reweighted ensemble densities.
**(C)** Mean JSD of within-state and between-state comparisons for APO, ADP and PEP states. Within-state comparisons (block diagonal) are consistently smaller than between-state comparisons, supporting robustness of feature selection.
**(D-F)** Same analysis as in (A-C), with the ensembles projected onto a TICA space (TIC0–TIC1) derived from BB torsion features.
**(G-I)** Same analysis as in (A-C), with the ensembles projected onto a TICA space (TIC0–TIC1) derived from HBOND-DIS features.

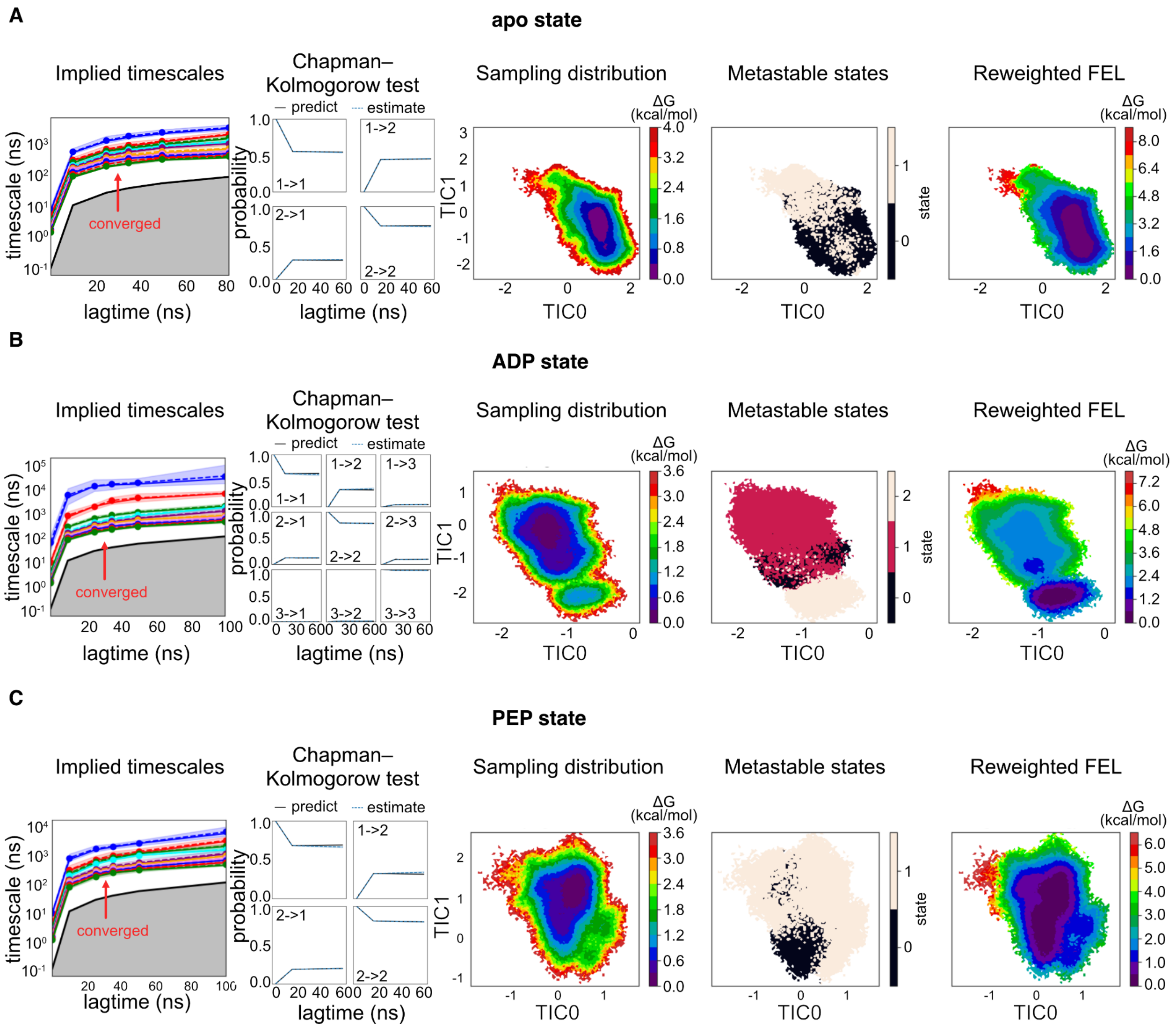


**Figure S7. MSM construction and validation for EcPFK, related to Figure 3.**

MSM construction and validation for **(A)** apo state, **(B)** ADP state, and **(C)** PEP state. From left to right, figure panels show the implied timescale plots with confidence intervals, Chapman-Kolmogorov tests of the Markov models, and comparisons between sampling density from raw trajectories, metastable states, together with the corresponding reweighted free energy landscapes (FELs).

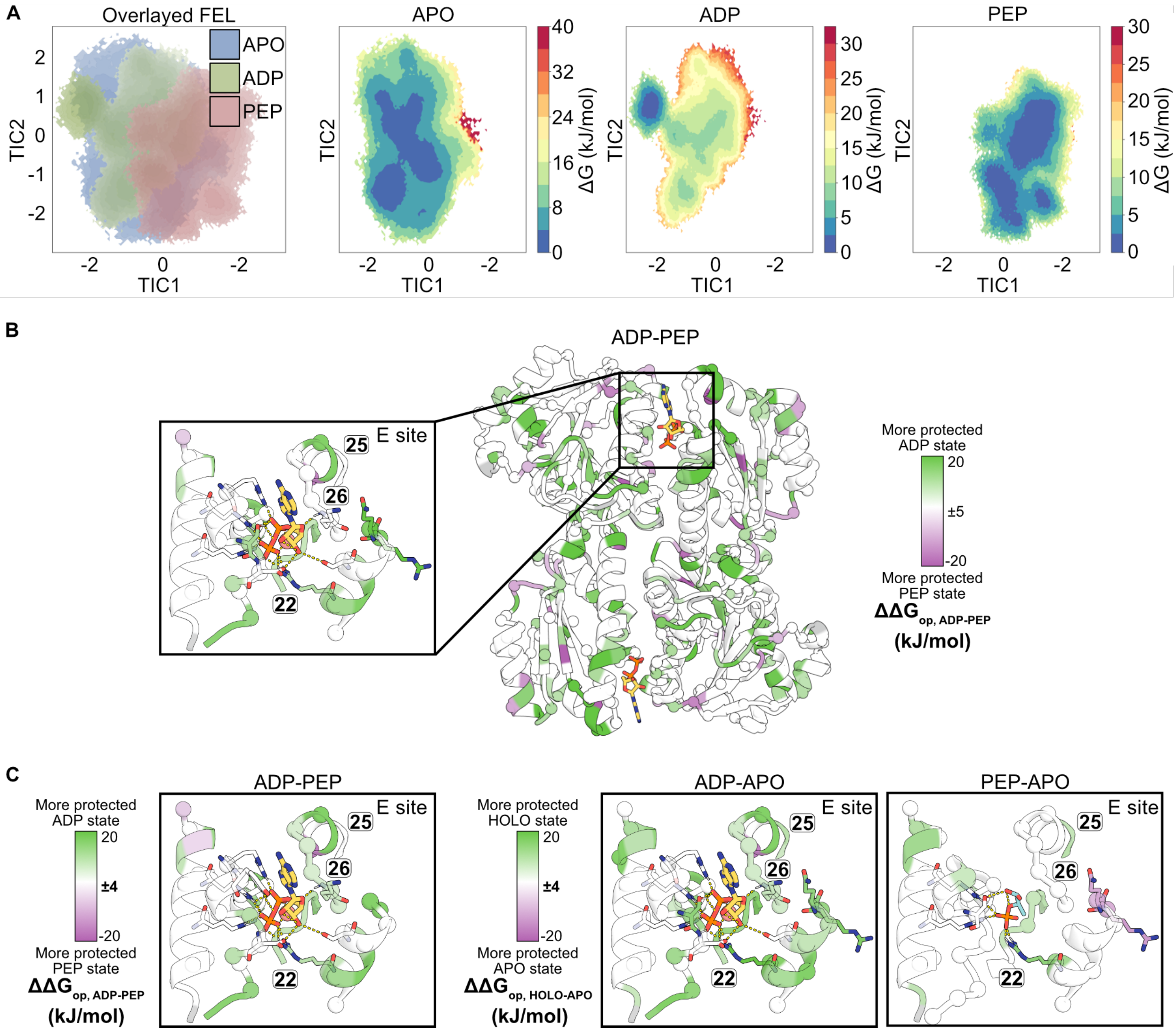


**Figure S8. Comparing MSMs with HX/MS-derived $\Delta G_{op}$, related to Figure 3.**

**(A)** FELs of the apo, ADP-, and PEP-bound ensembles projected onto TIC1 and TIC2.
**(B)** $\Delta\Delta G_{op,\ ADP\text{-}PEP}$ mapped onto the EcPFK crystal structure (PDB: 1PFK). Residues stabilized in the ADP-bound ensemble are green, while residues stabilized in the PEP-bound ensemble are purple. A threshold of ±5 kJ/mol is applied; residues with $-5 \leq \Delta G_{op} \leq 5$ kJ/mol are white.
**(C)** Zoomed E site for each binary state comparison, shown with a threshold of ±4 kJ/mol (residues with $-4 \leq \Delta G_{op} \leq 4$ kJ/mol are white).

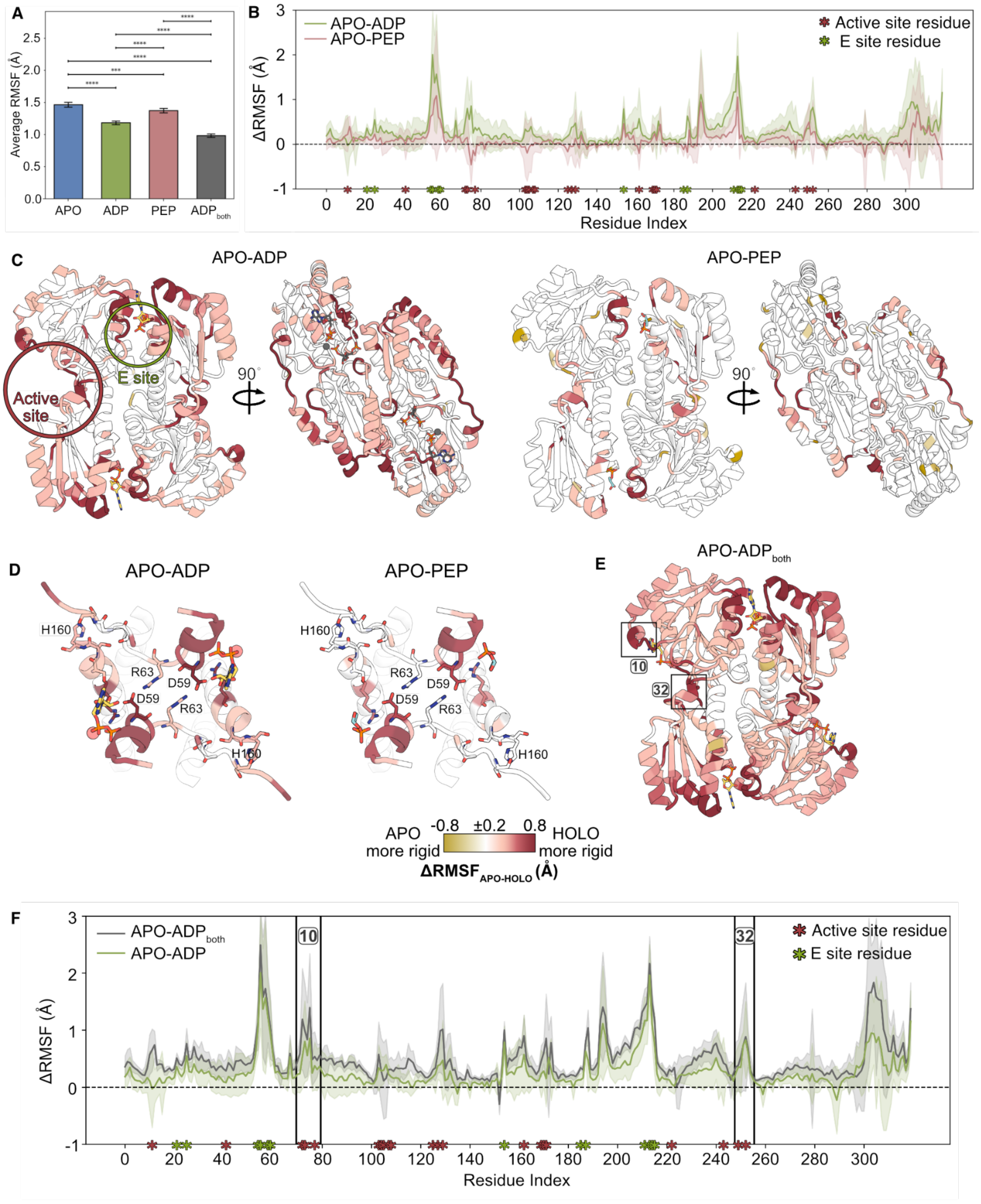


**Figure S9. RMSF analysis for EcPFK functional states, related to Figure 3.**

**(A)** Average heavy-atom RMSF for each ensemble. Error bars represent the 95% confidence interval. Asterisks denote statistical significance as determined by Welch's unpaired *t*-test (*p*<1e-4 ****, *p*<1e-3 ***,

*p*<0.01 **, *p*<0.05 *). $ADP_{both}$ refers to the PFK ensemble that has ADP bound in both the active and E sites.
**(B)** Difference in per-residue heavy-atom RMSF between the APO and ligand-bound ensembles ($\Delta RMSF_{APO\text{-}HOLO}$). Shaded regions indicate the 95% confidence interval.
**(C)** $\Delta RMSF_{APO\text{-}HOLO}$ mapped on the EcPFK dimer (PDB: 1PFK).
**(D)** Zoomed top-down view of the protomer-protomer interface showing ligand-dependent changes in flexibility.
**(E)** Per-residue $\Delta RMSF_{APO\text{-}HOLO}$ comparing ensembles with ADP bound in the E site only and with ADP bound in both the E and active sites. Segments 10 and 32, which are stabilized in both conditions, are labeled.
**(F)** EcPFK structure colored by ΔRMSF between the APO and ADP-bound ensembles (ADP in the E site only, and ADP in both the E and active sites), highlighting the positions of segments 10 and 32 within the active site.

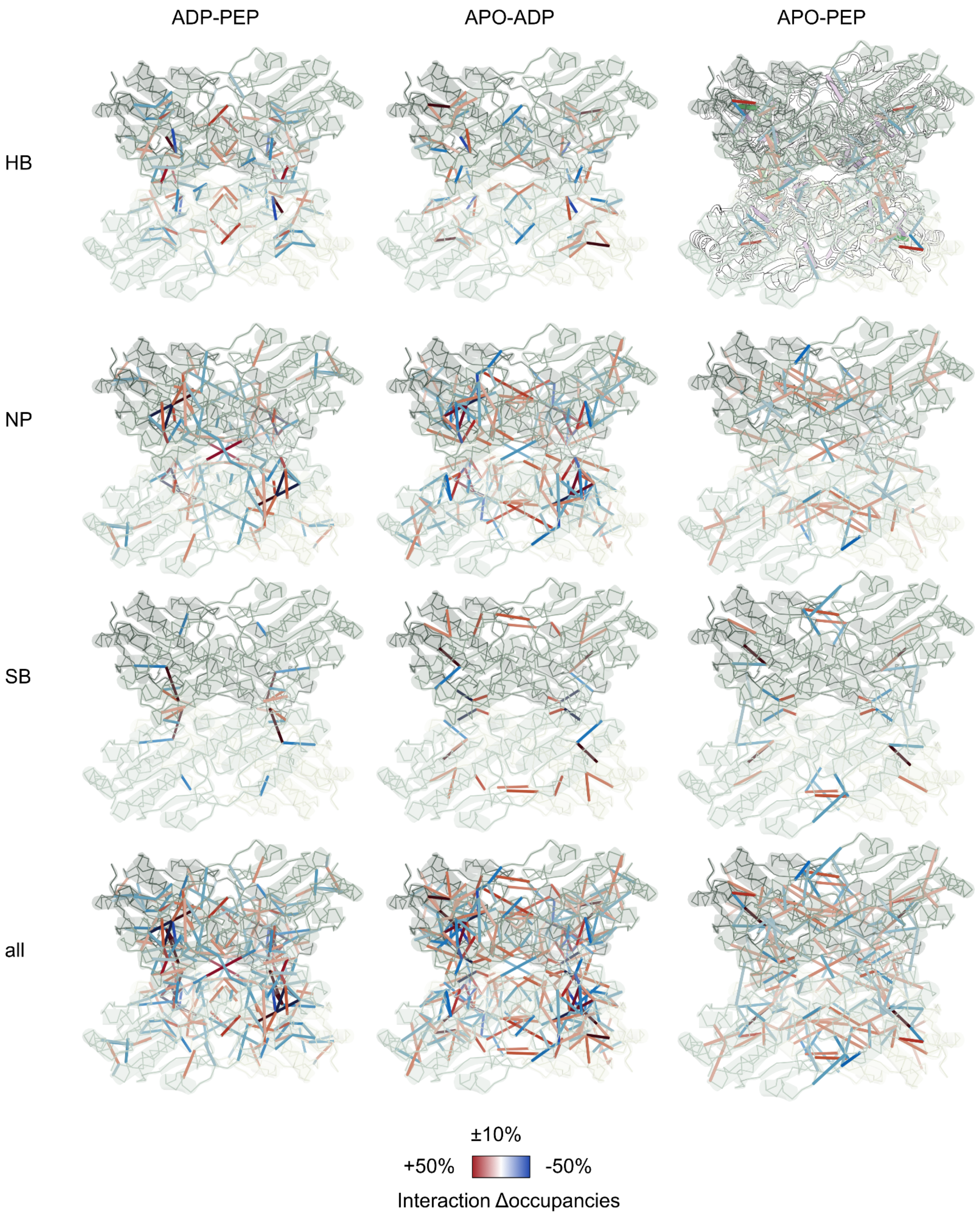


**Figure S10. Changes in interactions in reweighted ensembles, related to Figure 4.**

Pairwise interaction rearrangements across the apo, ADP, and PEP states. Three types of interactions are shown: hydrogen bonds (HB), nonpolar contacts (NP) and salt bridges (SB).

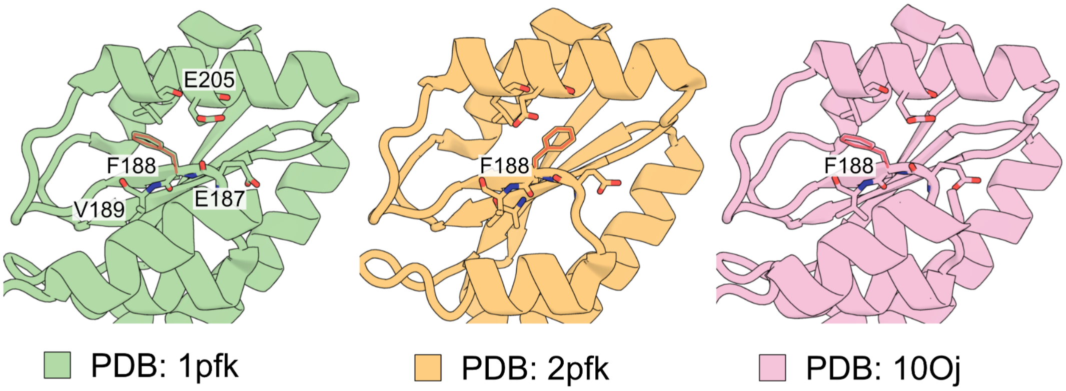


**Figure S11. F188 as a molecular switch, related to Figure 4.**

Alternative side-chain rotamers of F188 observed in the EcPFK crystal structures 1PFK and 2PFK (Rypniewski and Evans, 1989; Shirakihara and Evans, 1988) [3,6].

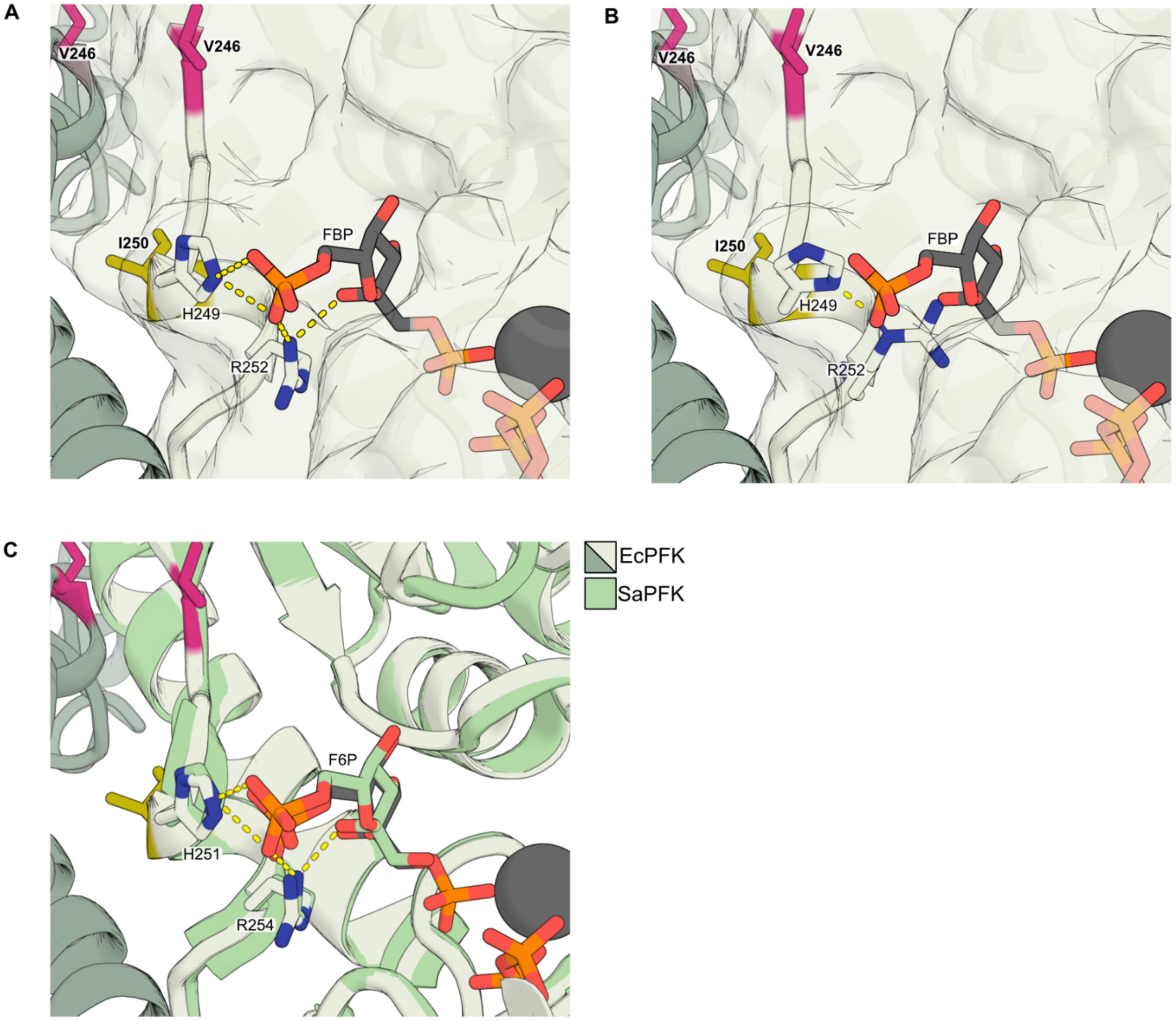


**Figure S12. Substrate binding in the PEP state, related to Figure 5.**

**(A)** Crystal structure of ADP-bound EcPFK (PDB: 1PFK) illustrating the binding mode of fructose-1,6-bisphosphate (FBP) within the substrate binding-pocket, with residues H249 and R252 shown as sticks.
**(B)** In the PEP-bound ensemble, the H249-R252 hydrogen bond exhibits increased occupancy relative to the ADP-bound ensemble. This residue configuration sterically clashes with the aligned FBP shown in (A) within the substrate-binding pocket. H249 and R252 rotamers are shown from a representative MD frame of the PEP-bound reweighted ensemble.
**(C)** Structural alignment of EcPFK (PDB: 1PFK) with fructose-6-phosphate (F6P)-bound *Staphylococcus aureus* PFK (PDB: 5XZ7) demonstrates that the binding mode of FBP resembles the binding mode of the substrate F6P (Tian *et al*., 2018) [7].

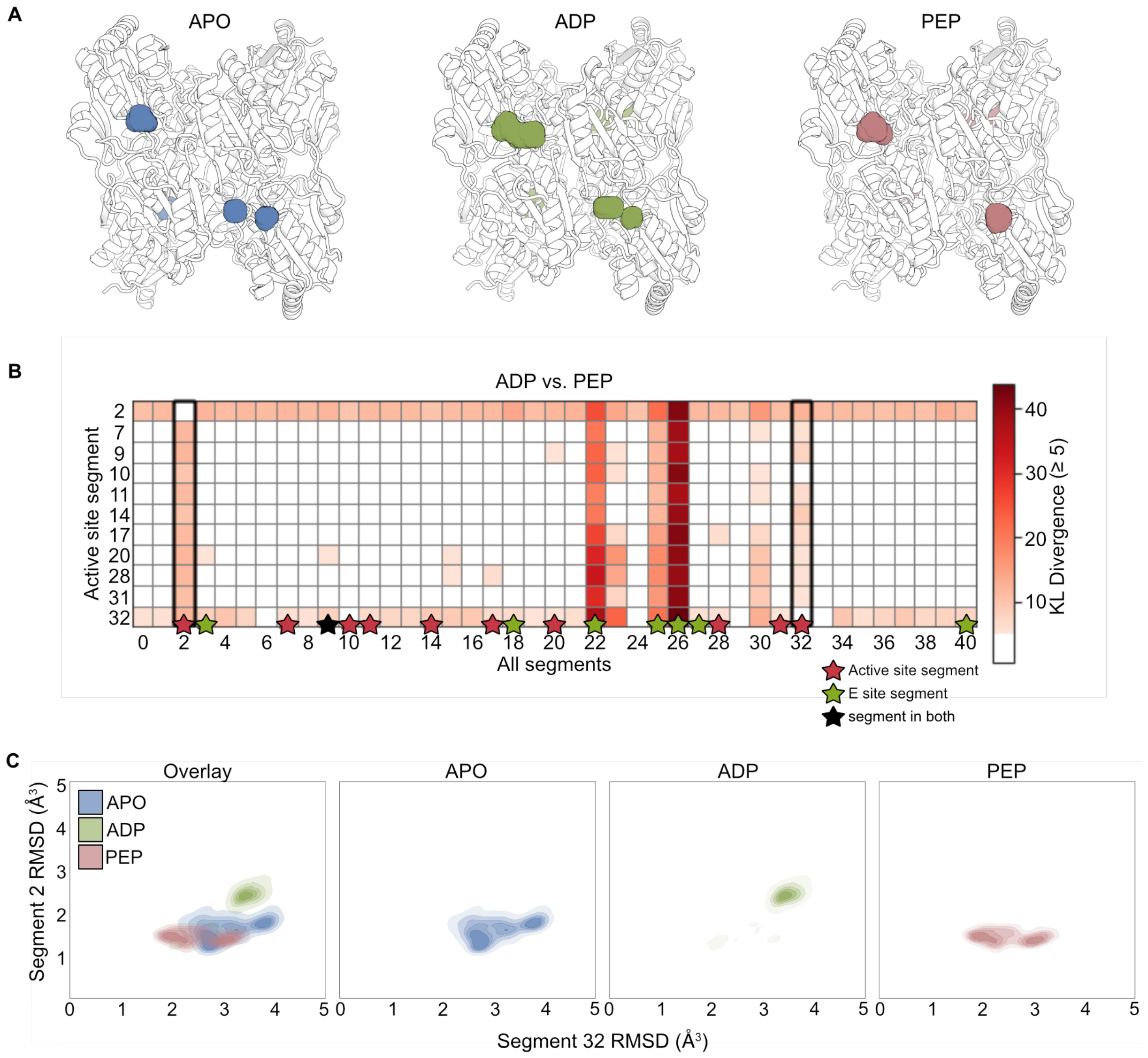


**Figure S13. Average active site volume for EcPFK functional states, related to Figure 5.**

**(A)** EcPFK MD simulation reference structures with spheres marking the active site cavity measured to calculate pocket volume in each ensemble.

**(B)** Kullback-Leibler (KL) divergence between secondary structure RMSD distributions of each active site segment and all other segments in EcPFK, computed for ADP vs. PEP. KL divergence identifies segments whose conformational responses to ADP vs. PEP differ in both direction and magnitude relative to other segments (*i.e.*, for each active site segment *i*, which segments *j* show the greatest divergence in ADP-PEP-dependent conformational reweighting relative to *i*?). While several non-active site segments (*e.g.*, segments 22 and 26) also exhibit high KL divergence with active site segments, segments 2 and 32 emerge as the most divergent within the active site, with conformational responses that are the most distinct relative to the remainder of the enzyme.

**(C)** Joint secondary structure RSMD distribution of segment 2 vs. segment 32. ADP binding stabilizes distinct conformations of segments 2 and 32 that are not sampled by the APO or PEP ensembles.

Location of segments 2 and 32 can be found in Fig. 2D. These segments contain several fructose binding residues.

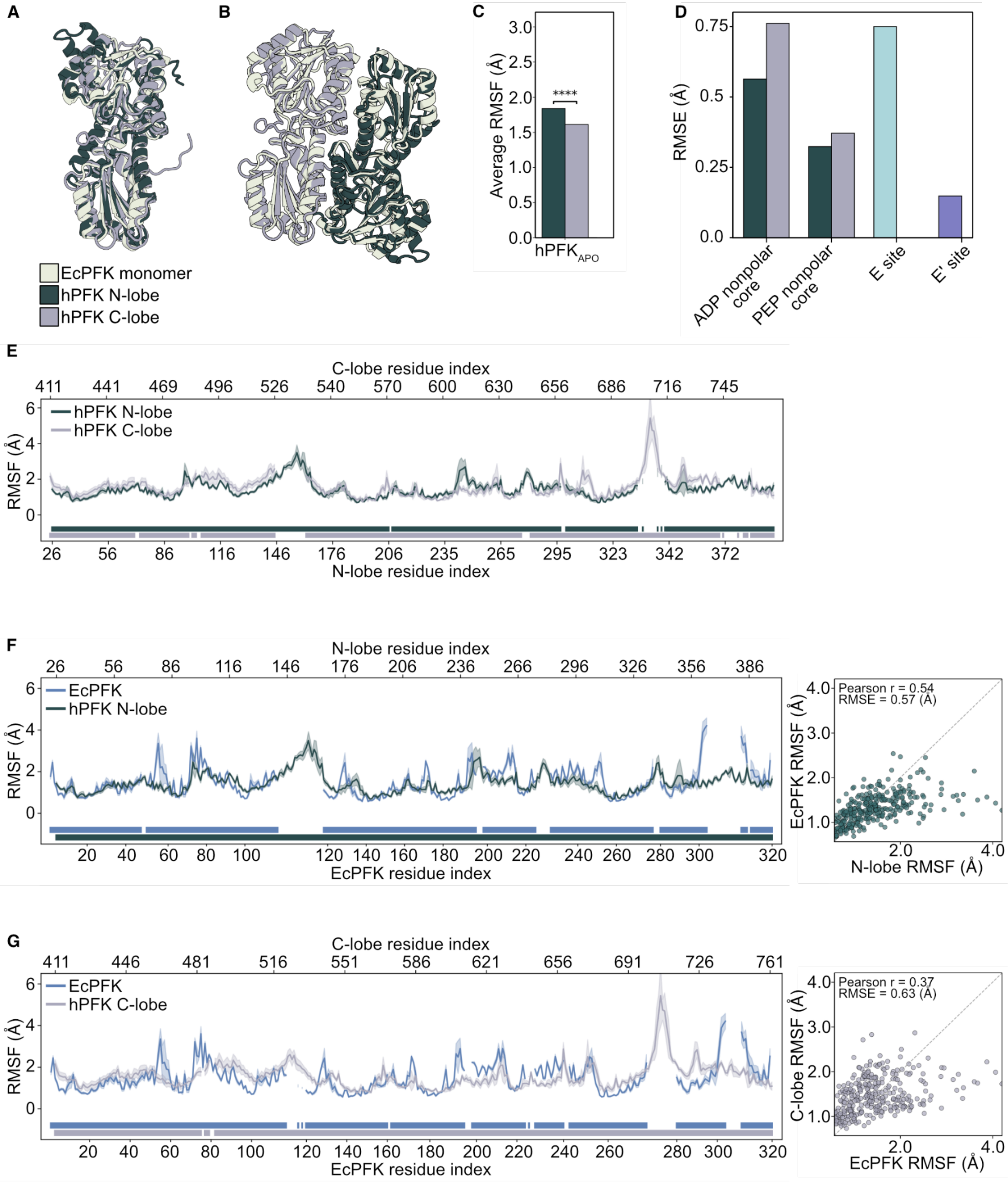


**Figure S14. Evolutionary origins of hPFK ligand regulation, related to Figure 6.**

**(A)** Structural alignment of the EcPFK monomer vs. the hPFK N- and C-terminal lobes. The ancestral and modern lobes in the hPFK monomer are differentiated by color (PDB: 1PFK (EcPFK); 4XYJ (hPFK)) (Shirakihara and Evans, 1988; Webb *et al.*, 2015) [6,8].

**(B)** Structural alignment of the EcPFK dimer and the hPFK monomer.
**(C)** Significant difference in average heavy-atom RMSF between the hPFK N- and C-terminal lobes. Statistical significance was assessed using Welch's *t*-test (**** $p < 1e\text{-}4$).
**(D)** Root mean square error (RMSE) calculated for aligned per-residue RMSF values between EcPFK and each hPFK lobe across various key regions. The E site and E' site are shown in distinct colors as each includes residues from both N- and C-terminal lobes. This panel accompanies Fig. 6D (See Table S3 for definitions of each site).
**(E)** Average per-residue heavy atom RMSF of aligned residues in the hPFK N- and C-terminal lobes. Shaded regions represent the 95% confidence interval. Rectangular bars beneath the curves indicate the sequence alignment between the two lobes.
**(F)** Average per-residue heavy atom RMSF of EcPFK from the apo ensemble plotted against aligned residues ($n$ = 316) of the hPFK N-terminal, and
**(G)** the C-terminal aligned residues ($n$ = 317).

**Supplemental tables**

**Table S1. HX experiment summary table.**

All experiments were performed with the KEEP setting in PIGEON (Lu, Wells *et al.*, 2026) [9]. Protease abbreviations: FP, immobilized protease type XIII. Nep, nepenthesin 2. AP, alanyl aminopeptidase.

| Dataset | **EcPFK Rep1** | **EcPFK Rep2** | **EcPFK Rep3** | **EcPFK Rep4** | **EcPFK Rep5** | **EcPFK Rep6** | **EcPFK Rep7** |
|---|---|---|---|---|---|---|---|
| Protein states | APO, ADP, PEP | APO, ADP, PEP | APO, ADP, PEP | APO, ADP, PEP | APO, ADP, PEP | APO, ADP, PEP | APO, ADP, PEP |
| Date | 04/24/23 | 07/07/23 | 08/29/23 | 07/07/23 | 08/29/23 | 01/21/25 | 01/22/25 |
| Protease column | FP | FP | FP | FP | FP | AP/pepsin | NP/pepsin |
| Reaction details | 50 mM HEPES, 100 mM KCl, 10 mM $MgCl_2$, 1 mM TCEP, pH 7.5, temperature 15 °C, final $D_2O$ concentration 90% | | | | | | |
| Time course (s) | 30, 120, 480 | 30, 120, 480, 1920, 7680 | 30, 120, 480, 960, 1920, 7680 | 30, 120, 480, 1920, 7680 | 30, 120, 480, 960, 1920, 7680 | 43, 342, 3041, 30043 | 100, 1030, 10030, 10531, 11030, 95861, 98985, 102105 |
| # of time points | 3 | 5 | 6 | 5 | 6 | 4 | 8 |
| Control samples | Maximally-labeled sample (full-D) | | | | | | |
| Back-exchange (mean / IQR) | 0.37/0.12 | 0.34/0.14 | 0.35/0.15 | 0.34/0.13 | 0.36/0.13 | 0.38/0.11 | 0.36/0.10 |
| # of peptides | 309 | 242 | 257 | 251 | 233 | 278 | 453 |
| Sequence coverage | 0.98 | 0.94 | 0.98 | 0.92 | 0.96 | 0.98 | 0.98 |
| Average peptide length / Redundancy | 1.1 | 1.5 | 1.4 | 1.4 | 1.6 | 1.2 | 0.7 |

**Table S2. PFNet results summary.**

The residue-level $\Delta G_{op}$ was determined by peptide coverage analysis as described previously (Lu, Wells, *et al.*, 2026) [9]. Prediction confidence is reported as a direct output of PFNet (Lu, Weber, *et al.*, 2024) [10], with $\Delta G_{op}$>0.8 classified as high-confidence. AE_mean and AE_median denote the mean and median isotopic mass envelope fitting errors, respectively, calculated between PFNet-reconstructed envelopes and the corresponding experimental envelopes.

| State | **EcPFK APO** | **EcPFK ADP** | **EcPFK PEP** |
|---|---|---|---|
| Highest log($k_{ex}$) | 0.65 | 0.35 | 0.53 |
| Lowest log($k_{ex}$) | -9.78 | -9.97 | -9.85 |
| Std log($k_{ex}$) | 3.05 | 3.15 | 3.05 |
| # single resolved residues | 251 | 261 | 258 |
| # non covered residues | 7 | 7 | 7 |
| # high-confidence residues (>0.8) | 204 (62.58%) | 162 (49.69%) | 190 (58.28%) |
| AE_mean_pfnet | 0.32 | 0.32 | 0.34 |
| AE_median_pfnet | 0.28 | 0.28 | 0.30 |

**Table S3. DSSP-defined secondary structure segment indices.**

The DSSP algorithm (Kabsch and Sander, 1983) [11] was used to assign secondary structure to individual residues in the ADP-bound EcPFK crystal structure (PDB: 1PFK) and to define contiguous secondary-structure segments, which were indexed and used consistently throughout this study.

| Segment index | Residue start | Residue end | DSSP | Residues |
|---|---|---|---|---|
| 0 | 0 | 2 | C | MET0- LYS2 |
| 1 | 3 | 8 | E | LYS3-THR8 |
| 2 | 9 | 15 | C | SER9-GLY15 |
| 3 | 16 | 29 | H | MET16-THR29 |
| 4 | 30 | 32 | C | GLU30-LEU32 |
| 5 | 33 | 37 | E | GLU33-ILE37 |
| 6 | 38 | 40 | C | TYR38- GLY40 |
| 7 | 41 | 46 | H | TYR41- GLU46 |
| 8 | 49 | 51 | E | MET49- GLN51 |
| 9 | 52 | 73 | C | LEU52-PHE73 |
| 10 | 74 | 77 | H | PRO74-ARG77 |
| 11 | 79 | 92 | H | GLU79-ARG92 |
| 12 | 93 | 95 | C | GLY93-ASP95 |
| 13 | 96 | 101 | E | ALA96-GLY101 |
| 14 | 103 | 113 | H | ASP103-THR113 |
| 15 | 114 | 118 | C | GLU114-PRO118 |
| 16 | 119 | 124 | E | CYS119-GLY124 |
| 17 | 125 | 136 | C | THR125-THR136 |
| 18 | 139 | 160 | H | PHE139-HIS160 |
| 19 | 163 | 168 | E | ILE163-VAL168 |
| 20 | 169 | 174 | C | MET169-GLY174 |
| 21 | 175 | 184 | H | ASP175-GLY184 |
| 22 | 185 | 187 | C | GLY185- GLU187 |
| 23 | 188 | 190 | E | PHE188-VAL190 |
| 24 | 191 | 197 | C | VAL191-SER197 |
| 25 | 198 | 211 | H | ARG198-LYS211 |
| 26 | 212 | 215 | C | GLY212-HIS215 |
| 27 | 216 | 221 | E | ALA216-THR221 |
| 28 | 222 | 226 | C | GLU222-ASP226 |

| | | | | |
|---|---|---|---|---|
| 29 | 227 | 238 | H | VAL227-THR238 |
| 30 | 239 | 241 | C | GLY239-GLU241 |
| 31 | 242 | 246 | E | THR242-VAL246 |
| 32 | 248 | 252 | H | GLY248-ARG252 |
| 33 | 253 | 257 | C | GLY253-VAL257 |
| 34 | 258 | 277 | H | PRO258-ALA277 |
| 35 | 278 | 281 | C | GLY278-GLY281 |
| 36 | 282 | 287 | E | ARG282-GLN287 |
| 37 | 290 | 295 | E | GLN290- ASP295 |
| 38 | 296 | 300 | H | ILE296-ILE300 |
| 39 | 301 | 308 | C | GLU301-LYS308 |
| 40 | 309 | 318 | H | GLY309-LEU318 |
| 41 | 319 | 319 | C | TYR319 |

**Table S4. Site definitions key functional areas in PFK, related to Figure 6.**

Residues used to calculate the Pearson correlation calculation between EcPFK ADP- and PEP-binding induced nonpolar cores and corresponding homologous residues in the N- and C-terminal lobes of hPFK for Fig. 6D and S14D.

| Site | EcPFK residue (PDB: 1PFK) | hPFK N-lobe residue (PDB: 4XYJ) | hPFK C-lobe residue (PDB: 4XYJ) |
|---|---|---|---|
| ADP nonpolar core | F188, F196, V189, L312, A181, V194 | W236, P244, V237, L391, A229, S242 | A602, F610, A603, R755, G595, E608 |
| PEP nonpolar core | I205, L230, A216, V218, I163, V165, F188 | L256, I288, N267, I269, T211, V213, W236 | L622, I650, G633, V635, V577, I579, A602 |

Residues used to calculate the Pearson correlation coefficient between EcPFK and hPFK for Fig. 6D and S14D. Residues included here are those listed in Shirakihara and Evans, 1988 [6] as residues involved in Mg-ADP binding in the E site of EcPFK.

| Site | EcPFK residue (PDB: 1PFK) | hPFK residue (PDB: 4XYJ) |
|---|---|---|
| E site | R21, R25, R54, Y55, S58, D59, **R154, G185, E187, K211, K214, H215, Y319** | M202, G233, D235, R262, K264, R265, L266, A398, **R430, R434, W463, T464, G467, G468** |
| E’ site | R21, R25, R54, Y55, S58, D59, **R154, G185, E187, K211, K214, H215, Y319** | R44, R48, W79, E80, S83, S84, **K567, G599, D601, T628, I630, Q631, R632, A762** |

Bold residues are those on the opposite protomer for EcPFK and residues on the C-terminal lobe for hPFK.

**Table S5. DNA sequence for plasmid used in this study.**

| Plasmid | DNA sequence |
|---|---|
| pET9a-6×His-TEV-EcPFK | TTCTTAGAAAAACTCATCGAGCATCAAATGAAACTGCAATTTATTCATATCAGGATTA<br>TCAATACCATATTTTTGAAAAAGCCGTTTCTGTAATGAAGGAGAAAACTCACCGAGGC<br>AGTTCCATAGGATGGCAAGATCCTGGTATCGGTCTGCGATTCCGACTCGTCCAACATC<br>AATACAACCTATTAATTTCCCCTCGTCAAAAATAAGGTTATCAAGTGAGAAATCACCA<br>TGAGTGACGACTGAATCCGGTGAGAATGGCAAAAGCTTATGCATTTCTTTCCAGACTT<br>GTTCAACAGGCCAGCCATTACGCTCGTCATCAAAATCACTCGCATCAACCAAACCGTT<br>ATTCATTCGTGATTGCGCCTGAGCGAGACGAAATACGCGATCGCTGTTAAAAGGACAA<br>TTACAAACAGGAATCGAATGCAACCGGCGCAGGAACACTGCCAGCGCATCAACAATAT<br>TTTCACCTGAATCAGGATATTCTTCTAATACCTGGAATGCTGTTTTCCCGGGGATCGC<br>AGTGGTGAGTAACCATGCATCATCAGGAGTACGGATAAAATGCTTGATGGTCGGAAGA<br>GGCATAAATTCCGTCAGCCAGTTTAGTCTGACCATCTCATCTGTAACATCATTGGCAA<br>CGCTACCTTTGCCATGTTTCAGAAACAACTCTGGCGCATCGGGCTTCCCATACAATCG<br>ATAGATTGTCGCACCTGATTGCCCGACATTATCGCGAGCCCATTTATACCCATATAAA<br>TCAGCATCCATGTTGGAATTTAATCGCGGCCTCGAGCAAGACGTTTCCCGTTGAATAT<br>GGCTCATAACACCCCTTGTATTACTGTTTATGTAAGCAGACAGTTTTATTGTTCATGA<br>CCAAAATCCCTTAACGTGAGTTTTCGTTCCACTGAGCGTCAGACCCCGTAGAAAAGAT<br>CAAAGGATCTTCTTGAGATCCTTTTTTTCTGCGCGTAATCTGCTGCTTGCAAACAAAA<br>AAACCACCGCTACCAGCGGTGGTTTGTTTGCCGGATCAAGAGCTACCAACTCTTTTTC<br>CGAAGGTAACTGGCTTCAGCAGAGCGCAGATACCAAATACTGTCCTTCTAGTGTAGCC<br>GTAGTTAGGCCACCACTTCAAGAACTCTGTAGCACCGCCTACATACCTCGCTCTGCTA<br>ATCCTGTTACCAGTGGCTGCTGCCAGTGGCGATAAGTCGTGTCTTACCGGGTTGGACT<br>CAAGACGATAGTTACCGGATAAGGCGCAGCGGTCGGGCTGAACGGGGGGTTCGTGCAC<br>ACAGCCCAGCTTGGAGCGAACGACCTACACCGAACTGAGATACCTACAGCGTGAGCTA<br>TGAGAAAGCGCCACGCTTCCCGAAGGGAGAAAGGCGGACAGGTATCCGGTAAGCGGCA<br>GGGTCGGAACAGGAGAGCGCACGAGGGAGCTTCCAGGGGGAAACGCCTGGTATCTTTA<br>TAGTCCTGTCGGGTTTCGCCACCTCTGACTTGAGCGTCGATTTTTGTGATGCTCGTCA<br>GGGGGGCGGAGCCTATGGAAAAACGCCAGCAACGCGGCCTTTTTACGGTTCCTGGCCT<br>TTTGCTGGCCTTTTGCTCACATGTTCTTTCCTGCGTTATCCCCTGATTCTGTGGATAA<br>CCGTATTACCGCCTTTGAGTGAGCTGATACCGCTCGCCGCAGCCGAACGACCGAGCGC<br>AGCGAGTCAGTGAGCGAGGAAGCGGAAGAGCGCCTGATGCGGTATTTTCTCCTTACGC<br>ATCTGTGCGGTATTTCACACCGCATATATGGTGCACTCTCAGTACAATCTGCTCTGAT<br>GCCGCATAGTTAAGCCAGTATACACTCCGCTATCGCTACGTGACTGGGTCATGGCTGC<br>GCCCCGACACCCGCCAACACCCGCTGACGCGCCCTGACGGGCTTGTCTGCTCCCGGCA<br>TCCGCTTACAGACAAGCTGTGACCGTCTCCGGGAGCTGCATGTGTCAGAGGTTTTCAC<br>CGTCATCACCGAAACGCGCGAGGCAGCTGCGGTAAAGCTCATCAGCGTGGTCGTGAAG<br>CGATTCACAGATGTCTGCCTGTTCATCCGCGTCCAGCTCGTTGAGTTTCTCCAGAAGC<br>GTTAATGTCTGGCTTCTGATAAAGCGGGCCATGTTAAGGGCGGTTTTTTCCTGTTTGG<br>TCACTGATGCCTCCGTGTAAGGGGGATTTCTGTTCATGGGGGTAATGATACCGATGAA<br>ACGAGAGAGGATGCTCACGATACGGGTTACTGATGATGAACATGCCCGGTTACTGGAA<br>CGTTGTGAGGGTAAACAACTGGCGGTATGGATGCGGCGGGACCAGAGAAAAATCACTC<br>AGGGTCAATGCCAGCGCTTCGTTAATACAGATGTAGGTGTTCCACAGGGTAGCCAGCA<br>GCATCCTGCGATGCAGATCCGGAACATAATGGTGCAGGGCGCTGACTTCCGCGTTTCC<br>AGACTTTACGAAACACGGAAACCGAAGACCATTCATGTTGTTGCTCAGGTCGCAGACG<br>TTTTGCAGCAGCAGTCGCTTCACGTTCGCTCGCGTATCGGTGATTCATTCTGCTAACC<br>AGTAAGGCAACCCCGCCAGCCTAGCCGGGTCCTCAACGACAGGAGCACGATCATGCGC<br>ACCCGTGGCCAGGACCCAACGCTGCCCGAGATGCGCCGCGTGCGGCTGCTGGAGATGG<br>CGGACGCGATGGATATGTTCTGCCAAGGGTTGGTTTGCGCATTCACAGTTCTCCGCAA<br>GAATTGATTGGCTCCAATTCTTGGAGTGGTGAATCCGTTAGCGAGGTGCCGCCGGCTT<br>CCATTCAGGTCGAGGTGGCCCGGCTCCATGCACCGCGACGCAACGCGGGGAGGCAGAC<br>AAGGTATAGGGCGGCGCCTACAATCCATGCCAACCCGTTCCATGTGCTCGCCGAGGCG<br>GCATAAATCGCCGTGACGATCAGCGGTCCAGTGATCGAAGTTAGGCTGGTAAGAGCCG<br>CGAGCGATCCTTGAAGCTGTCCCTGATGGTCGTCATCTACCTGCCTGGACAGCATGGC |

| | |
|---|---|
| | CTGCAACGCGGGCATCCCGATGCCGCCGGAAGCGAGAAGAATCATAATGGGGAAGGCC<br>ATCCAGCCTCGCGTCGCGAACGCCAGCAAGACGTAGCCCAGCGCGTCGGCCGCCATGC<br>CGGCGATAATGGCCTGCTTCTCGCCGAAACGTTTGGTGGCGGGACCAGTGACGAAGGC<br>TTGAGCGAGGGCGTGCAAGATTCCGAATACCGCAAGCGACAGGCCGATCATCGTCGCG<br>CTCCAGCGAAAGCGGTCCTCGCCGAAAATGACCCAGAGCGCTGCCGGCACCTGTCCTA<br>CGAGTTGCATGATAAAGAAGACAGTCATAAGTGCGGCGACGATAGTCATGCCCCGCGC<br>CCACCGGAAGGAGCTGACTGGGTTGAAGGCTCTCAAGGGCATCGGTCGACGCTCTCCC<br>TTATGCGACTCCTGCATTAGGAAGCAGCCCAGTAGTAGGTTGAGGCCGTTGAGCACCG<br>CCGCCGCAAGGAATGGTGCATGCAAGGAGATGGCGCCCAACAGTCCCCCGGCCACGGG<br>GCCTGCCACCATACCCACGCCGAAACAAGCGCTCATGAGCCCGAAGTGGCGAGCCCGA<br>TCTTCCCCATCGGTGATGTCGGCGATATAGGCGCCAGCAACCGCACCTGTGGCGCCGG<br>TGATGCCGGCCACGATGCGTCCGGCGTAGAGGATCGAGATCTCGATCCCGCGAAATTA<br>ATACGACTCACTATAGGCCCCTCTAGAAATAATTTTGTTTAACTTTAAGAAGGAGATA<br>TACCCATGCATCACCATCACCATCACGAAAACTTATATTTCCAATCTATTAAGAAAAT<br>CGGTGTGTTGACAAGCGGCGGTGATGCGCCAGGCATGAACGCCGCAATTCGCGGGGTT<br>GTTCGTTCTGCGCTGACAGAAGGTCTGGAAGTAATGGGTATTTATGACGGCTATCTGG<br>GTCTGTATGAAGACCGTATGGTACAGCTAGACCGTTACAGCGTGTCTGACATGATCAA<br>CCGTGGCGGTACGTTCCTCGGTTCTGCGCGTTTCCCGGAATTCCGCGACGAGAACATC<br>CGCGCCGTGGCTATCGAAAACCTGAAAAAACGTGGTATCGACGCGCTGGTGGTTATCG<br>GCGGTGACGGTTCCTACATGGGTGCAATGCGTCTGACCGAAATGGGCTTCCCGTGCAT<br>CGGTCTGCCGGGCACTATCGACAACGACATCAAAGGCACTGACTACACTATCGGTTTC<br>TTCACTGCGCTGAGCACCGTTGTAGAAGCGATCGACCGTCTGCGTGACACCTCTTCTT<br>CTCACCAGCGTATTTCCGTGGTGGAAGTGATGGGCCGTTATTGTGGAGATCTGACGTT<br>GGCTGCGGCCATTGCCGGTGGCTGTGAATTCGTTGTGGTTCCGGAAGTTGAATTCAGC<br>CGTGAAGACCTGGTAAACGAAATCAAAGCGGGTATCGCGAAAGGTAAAAAACACGCGA<br>TCGTGGCGATTACCGAACATATGTGTGATGTTGACGAACTGGCGCATTTCATCGAGAA<br>AGAAACCGGTCGTGAAACCCGCGCAACTGTGCTGGGCCACATCCAGCGCGGTGGTTCT<br>CCGGTGCCTTACGACCGTATTCTGGCTTCCCGTATGGGCGCTTACGCTATCGATCTGC<br>TGCTGGCAGGTTACGGCGGTCGTTGTGTAGGTATCCAGAACGAACAGCTGGTTCACCA<br>CGACATCATCGACGCTATCGAAAACATGAAGCGTCCGTTCAAAGGTGACTGGCTGGAC<br>TGCGCCGAAAAAATGTATTAAGAGCTCCGTCGACAAGCTTGCGGCCGCACTCGAGCAC<br>CACCACCACCACCACTGAGATCCGGCTGCTAACAAAGCCCGAAAGGAAGCTGAGTTGG<br>CTGCTGCCACCGCTGAGCAATAACTAGCATAACCCCTTGGGGCCTCTAAACGGGTCTT<br>GAGGGGTTTTTTGCTGAAAGGAGGAACTATATCCGGATATCCACAGGACGGGTGTGGT<br>CGCCATGATCGCGTAGTCGATAGTGGCTCCAAGTAGCGAAGCGAGCAGGACTGGGCGG<br>CGGCCAAAGCGGTCGGACAGTGCTCCGAGAACGGGTGCGCATAGAAATTGCATCAACG<br>CATATAGCGCTAGCAGCACGCCATAGTGACTGGCGATGCTGTCGGAATGGACGATATC<br>CCGCAAGAGGCCCGGCAGTACCGGCATAACCAAGCCTATGCCTACAGCATCCAGGGTG<br>ACGGTGCCGAGGATGACGATGAGCGCATTGTTAGATTTCATACACGGTGCCTGACTGC<br>GTTAGCAATTTAACTGTGATAAACTACCGCATTAAAGCTTATCGATGATAAGCTGTCA<br>AACATGAGAA |

**Table S6. X-ray crystallography summary statistics.**

| **Data statistics for PBD 10OJ** | |
|---|---|
| Beamline | NSLS-II BEAMLINE 19-ID |
| Wavelength (Å) | 0.9795 |
| Space group | P 21 2 21 |
| a, b, c (Å) | 76.70 90.85 103.71 |
| α, β, γ (º) | 90.0 90.0 90.0 |
| $Z_a$[a] / Solvent content (%) | 49.0 |
| Resolution range (Å) | 45.42– 2.38 (2.47-2.36) |
| Total reflections | 396647 (38505) |
| Unique reflections | 29760 (3088) |
| Completeness (%) | 100 (100.0) |
| Multiplicity | 13.3 (12.5) |
| $CC_{1/2}$ | 0.995 (0.123) |
| I/σ | 7.0 (0.4) |
| $R_{merge}$ | 0.366 (6.692) |
| $R_{pim}$ | 0.148 (2.039) |
| **Refinement statistics** | |
| Reflection used | 29302 (3979) |
| R-factor | 0.247 |
| Free R-factor | 0.279 |
| RMSD from ideal: | |
| Bond length (Å) | 0.005 |
| Angle (°) | 0.622 |
| Ramachandran (%): | |
| Favored | 96.23 |
| Allowed | 3.45 |

[a] Za stands for number of subunits per asymmetric unit.

### Supplemental references

1. Evans, P.R., and Hudson, P.J. (1979). Structure and control of phosphofructokinase from Bacillus stearothermophilus. Nature *279*, 500–504.

2. Schirmer, T., and Evans, P.R. (1990). Structural basis of the allosteric behaviour of phosphofructokinase. Nature *343*, 140–145.

3. Rypniewski, W.R., and Evans, P.R. (1989). Crystal structure of unliganded phosphofructokinase from Escherichia coli. Journal of Molecular Biology *207*, 805–821.

4. Prinz, J.-H., Wu, H., Sarich, M., Keller, B., Senne, M., Held, M., Chodera, J.D., Schütte, C., and Noé, F. (2011). Markov models of molecular kinetics: Generation and validation. The Journal of Chemical Physics *134*, 174105.

5. Endres, D.M., and Schindelin, J.E. (2003). A new metric for probability distributions. IEEE Transactions on Information Theory *49*, 1858–1860.

6. Shirakihara, Y., and Evans, P.R. (1988). Crystal structure of the complex of phosphofructokinase from Escherichia coli with its reaction products. Journal of Molecular Biology *204*, 973–994.

7. Tian, T., Wang, C., Wu, M., Zhang, X., and Zang, J. (2018). Structural Insights into the Regulation of *Staphylococcus aureus* Phosphofructokinase by Tetramer–Dimer Conversion. Biochemistry *57*, 4252–4262.

8. Webb, B.A., Forouhar, F., Szu, F.-E., Seetharaman, J., Tong, L., and Barber, D.L. (2015). Structures of human phosphofructokinase-1 and atomic basis of cancer-associated mutations. Nature *523*, 111–114.

9. Lu, C., Wells, M.L., Reckers, A., McBride, S.K., and Glasgow, A. (2026). Site-resolved energetic information from HX–MS experiments. Nat. Chem. Biol. *22*, 307–317.

10. Lu, C., Weber, K.C., McBride, S.K., Reckers, A., and Glasgow, A. A machine learning method for calculating highly localized protein stabilities. BioRxiv Rev., 2025.10.21.683809.

11. Kabsch, W., and Sander, C. (1983). Dictionary of protein secondary structure: pattern recognition of hydrogen-bonded and geometrical features. PubMed.
